# Predicting gene sequences with AI to study codon usage patterns

**DOI:** 10.1101/2024.02.11.579798

**Authors:** Tomer Sidi, Shir Bahiri-Elitzur, Tamir Tuller, Rachel Kolodny

## Abstract

Selective pressure acts on the codon use, optimizing multiple, overlapping signals that are only partially understood. We trained artificial intelligence (AI) models to predict the codons given their amino acid sequence in the eukaryotes *Saccharomyces cerevisiae* and *Schizosaccharomyces pombe* and the bacteria *Escherichia coli* and *Bacillus subtilis*, to study the extent to which we can learn patterns in naturally occurring codons to improve predictions. We trained our models on a subset of the proteins, and evaluated their predictions on large, separate sets of proteins of varying lengths and expression levels. Our models significantly outperformed naïve frequency-based approaches, demonstrating that there are dependencies between codons that can be learned to better predict evolutionary-selected codon usage. The prediction accuracy advantage of our models is greater for highly expressed genes and it is greater in bacteria than eukaryotes, supporting the hypothesis that there is a monotonic relationship between selective pressure for complex codon patterns and effective population size. Also, in *S*. *cerevisiae* and bacteria, our models were more accurate for longer proteins, suggesting that the AI system may have learned patterns related to co-translational folding. Gene functionality and conservation were also important determinants that affect the performance of our models. Finally, we showed that using information encoded in homologous proteins has only a minor effect on prediction accuracy, perhaps due to complex codon-usage codes in genes undergoing rapid evolution. In summary, our study employing contemporary AI methods offers a new perspective on codon usage patterns and a novel tool to optimize codon usage in endogenous and heterologous proteins.

**Significance statement:** Can one predict codon sequences used by an organism to encode a given amino acid sequence? This is difficult, because there are exponentially many codon sequences that can encode the same amino acid sequence and evolution is stochastic. Indeed, codons frequencies vary, a phenomenon known as codon-bias, yet we improve upon frequency-based predictions using contemporary AI tools that learn complex patterns and capture interactions between codons. Because our predictions are tested fairly, on cases not seen during the training process, accurate predictions suggest that these learned patterns are not random, and may be related to the evolutionary process. Thus, studying where our predictions are more accurate, is expected to reveal novel insights related to the way evolution shapes coding regions.

## Introduction

Although in all known organisms, there are 61 codons that each encode one of the 20 amino acids, 18 amino acids are encoded by multiple (two to six) codons. Thus, there are many codon sequences that encode the same amino acid sequence. The selection of codons used impacts protein production, indirectly, by influencing the availability of tRNA and free ribosomes in the cell [1–4], and directly by influencing mRNA structure and stability [3, 5], transcription [6], splicing [7], and translation kinetics [8–10]. This translation kinetics, in turn, influence protein co-translational folding and regulation [9, 11, 12].

Certain codons are preferred over others, a phenomenon known as codon bias [1, 11], and this bias differs not only across species [9] but also within species in a manner that depends on expression level and protein length [5, 11, 13]. Codon usage even differs along individual genes. For example, there are different usage patterns at the beginning of coding regions versus protein domain boundaries [14, 15]. Use of rare codons can slow translation [11, 16] and enable the generation of functionally or structurally stable proteins [3]. Because codon usage influences efficiency and accuracy of protein synthesis, it was recognized as a code within the genetic code that is subject to evolutionary selection [3, 13].

Computationally predicting the codon encodings of proteins in different organisms holds practical value, even in studies like ours that do not emulate the evolutionary process. Codon usage can have a significant effect on protein levels in different organisms [17–20]. Production of heterologous (non-host) proteins for use in protein science and biotechnology [21], in bacterial cell factories [22], as vaccines [23], or for agricultural purposes [24] requires optimization of coding sequence. The disparities in codon biases between the original and host organisms necessitate adjusting the codons to the biases of the new host. Indeed, dozens of measures aim to model codon usage patterns (reviewed in [25]). These measures, however, are usually limited to capturing local statistics of codon distribution.

To rigorously approach this prediction challenge, one should establish distinct training and test sets [26]. Two meaningful baseline models that were used in previous studies are the naïve Bayes predictor, which predicts the most frequently used codon for each amino acid [27] and the bigram frequency model which predicts the most frequently used codon, conditioned on its preceding codon [28, 29]. These frequencies can be estimated from the training set. There may be patterns of interactions among codons, that are more distant in primary sequence, and learning these may yield even better predictions [3, 30]. To learn codon patterns with the limited data available per organism, sophisticated tools that learn data distributions are needed [31].

Knowing the codons used in an orthologous protein in another organism may aid in prediction. For example, orthologous proteins may have a codon at a particular position, that differs due to usage bias, but that functions to induce translation pausing that is crucial for proper protein folding. If codons with unique functions can be identified and mimicked, the prediction may be more accurate [8, 22]. In support of the utility of this information, it was shown that position-dependent clusters of optimal and nonoptimal codons are conserved among orthologous proteins [32, 33], and that codon usage relates to protein structure with nonoptimal codons aligning domain boundaries [8, 9, 11, 34]. Indeed, these observations underlie the design of codon harmonization tools that predict a codon sequence for optimal incorporation of a given amino acid sequence in a non-native organism. Codon harmonization tools are not, however, designed for orthologous proteins but rather for those with the same amino acid sequence [35, 36]. In other words, they are not intended to predict the evolutionarily selected codons in one organism based on the codons of a different, albeit orthologous, protein. In practical terms, this means that codon harmonization tools mimic readily described properties like frequency rank [35, 36].

Deep networks, and transformers in particular [31], have emerged as the tool of choice to learn complex distributions like those characterizing codon usage. In prior research, deep networks have been used to address challenges in this domain: Most closely related is the work by Yang et al. who used deep networks (BiLSTM) to predict codon sequences of highly expressed proteins, but their predictions failed to improve upon the frequency-based baseline [26]. Other studies suggest however that there is a signal within codon sequences that deep networks can harvest: Tunney et al. used a feed-forward network to successfully predict the ribosome density from the sequence neighborhood [37], and several studies have demonstrated the utility of deep networks (RNN, T5, and BiLSTM) to optimize the gene expression levels of two to four proteins [38–40], and predict other aspects of gene expression [41–44].

Here, we take a data-driven approach to investigate the naturally occuring codon sequences in four organisms: the eukaryotes *S. cerevisiae* and *S. pombe*, and the bacteria *E. coli* and *B. subtilis*. We explored two scenarios: predicting the codon sequence from amino acid sequences, and predicting the codon sequence in one organism given the codon sequence of an orthologous protein in another organism. We used mBART, a transformer-based encoder-decoder architecture that extends BART, developed for a single natural language [45, 46]. mBART learns a shared model for multiple languages (e.g., English and French) allowing it both generate text in these languages and convert (i.e., translate) text from one language to another. In our setting, the analog of a language is an organism, and our mBART-trained models can both generate codon sequences for multiple organisms and mimic the codons of an orthologous protein. That our best models outperformed the frequency-based baselines, suggesting that there are patterns of codon interactions between residues (including non-neighboring residues) that can be learned. The ability of the AI model to learn such patterns, and specifically how much it improves the prediction differs as a function of expression level and protein length in different organisms, and suggests where the complex patterns in naturally occurring proteins are selected for. Furthermore, in *S. cerevisiae* and *E. coli*, we compared the accuracy gain of our model to the frequency-based baseline for functional sets of proteins grouped by Gene Ontology (GO) annotations and found that accuracy gain was higher than average for some molecular functions and biological processes. The novel artificial intelligence (AI) tool introduced here, with publicly available code and an easy-to-use web interface (https://www.aa2codons.info/), will enable future investigations related to evolutionary selection of codon sequences.

## Results

### (1) Training of mBART models to predict codons from amino acid sequence using masking and mimicking

We consider two tasks: masking, which is prediction of codons from the amino acid sequence, and mimicking, which is prediction of codons based on codons of an ortholog protein in another organism. The rationale for the mimicking mode is that the rate of translation elongation depends on the codon used [47], and the nonuniform rate may be important for co-translational protein folding [8, 48, 49]. Thus, as codons of orthologous proteins in two organisms may encode similar elongation rates, the codon sequence of the orthologous protein may be useful in prediction of codon usage for a protein of interest.

We trained several mBART models [45, 46] to support masking and mimicking tasks in four well studied model organisms: *S. cerevisiae, S. pombe., E. coli,* and *B. subtilis.* Following standard machine learning practices, we divided the protein data from these organisms into three distinct sets: ∼70% in the training set, ∼10% in the validation set, and ∼20% in the test set. All three sets included proteins with a wide range of expression levels. At the amino acid level, none of the proteins in the test set were closely related to those in the training set (based on amino-acid sequence clustering using CD-HIT [50] with a threshold of 0.7). The test set included 1240 *S. cerevisiae*, 1024 *S. pombe*, 812 *E. coli*, and 855 *B. subtilis* proteins (of which all but 496, 463, 271, and 247, respectively, have measured expression levels). Thus, evaluation of the models was conducted under stringent conditions, both in that the test set included a significant number of proteins with a wide range of expression levels and lengths and that the training set was not similar at the amino acid level to proteins in the test set (see Methods for more detail).

Figure 1 illustrates the training procedure and the input format of the mBART models. In masking mode, the input data is (only) the amino acid sequence of the target protein; in mimicking mode, the input data is the amino acid sequence *and* the codons of an ortholog protein (Figure 1A). The input format with two concatenated sequences supports both tasks and includes tokens indicating the organisms of the proteins (Figure 1B). We trained multiple models, each with a specific window size (Figure 1C). Pre-processing and post-processing steps are illustrated in Figure 1D. The performance of the trained models was evaluated based on accuracy of prediction of the codons of all proteins in the test set. We observed that the frequency-based model trained on highly expressed proteins was more accurate when predicting the codons of highly expressed proteins (Figure 2); therefore, we added a 6-class classification token of the expression level of the protein within its organism (omitted from the illustration in Figure 1A for brevity). In masking mode, the input is two copies of the same amino acid sequence and the organism token. In mimicking mode, the input is a gap-infused alignment of two orthologous proteins, where the first sequence is the codons of the source protein and its organism, and the second sequence is the amino acids of the target protein and its organism. Some of the codons in the target sequences may be passed as input for context (i.e., they are not predicted). For example, during pre-training, only 30% of the positions are masked by amino acids and predicted by the model.

**Figure 1:**
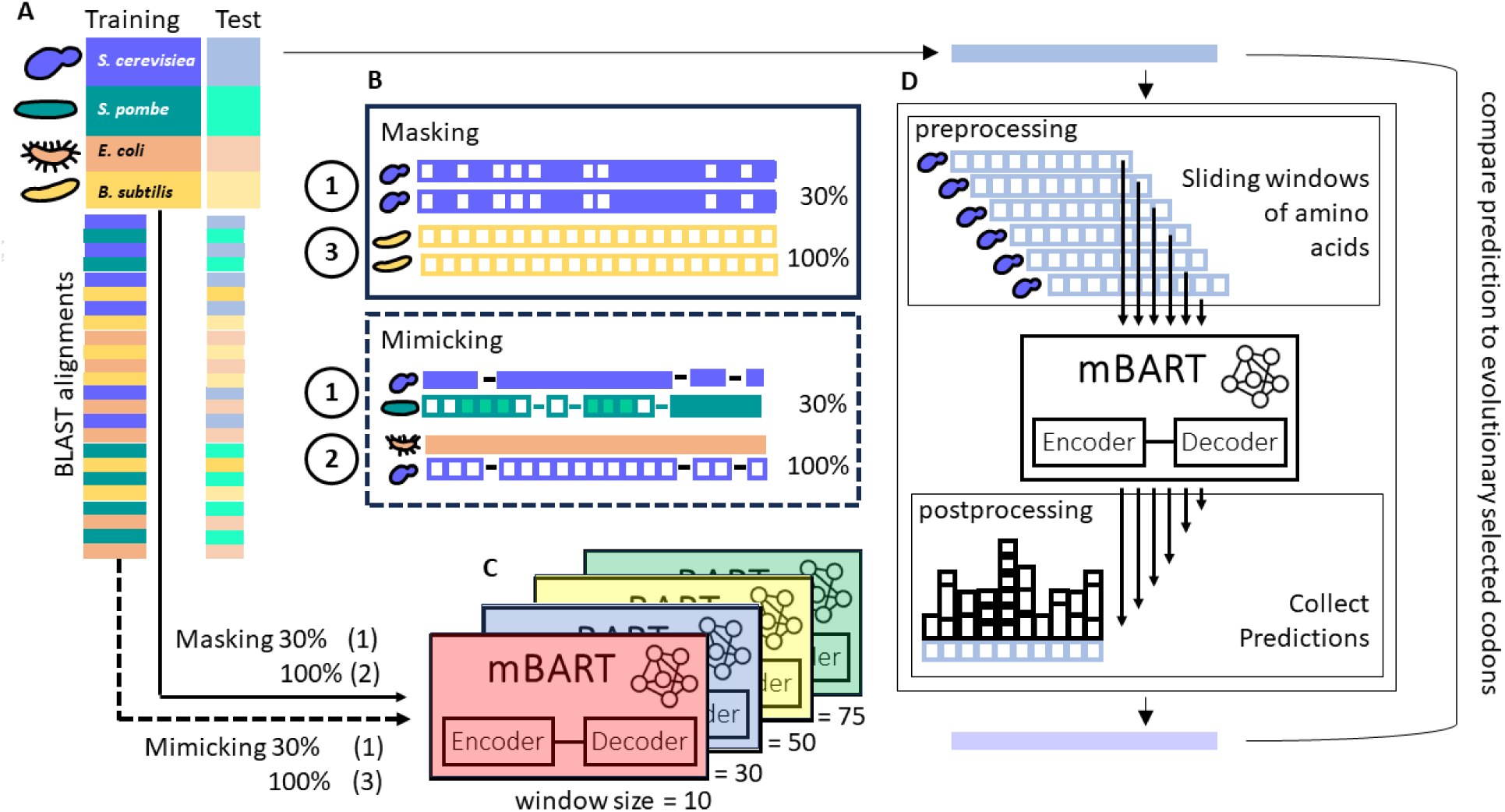
Strategy to learn codon usage patterns in *S. cerevisiae, S. pombe, E. coli*, and *B. subtilis*. (A) Our dataset includes protein amino sequence data from the four organisms and BLAST-identified protein alignments. Following standard practices in AI, we split our data into training and test sets (after clustering the proteins with CD-HIT and ensuring that closely related proteins were in either the training or the test set; see Methods for details). (B) To support both the masking mode and the mimicking mode, the input format has two sequences, with each sequence preceded by its source organism. In the cartoon representation, codons are shown by solid-colored boxes, and their corresponding amino acids by hollow boxes. In masking mode, both sequences are the same, and the input is either a sequence of codons where 30% or 100% of the positions are masked. In mimicking mode, the first sequence are the codons in an orthologous protein, and the second sequence is a masked sequence of codons, where 30% or 100% of the positions are masked. (C) The training set was used to train mBART models with varying window sizes (10, 30, 50, and 75 codons). First (1), we pre-trained the models with 30% masking and mimicking. Next (2), we fine-tuned the model with the 100% masked sequences to generate the fine-tuned masking model. Third (3), we further fine-tuned the model with 100% masking and mimicking data to generate the fine-tuned mimicking model. (D) During inference, there are pre- and post-processing steps: For each protein in the test set, all sliding windows corresponding to the model window size were considered, and each sequence of codons was fully masked. The cartoon example shows predictions in masking mode for a sliding window of 10 codons in an *S. cerevisiae* protein. These predictions are combined to yield the final codon prediction for the sequence, and we measured the accuracy of the prediction with respect to the evolutionarily selected codon sequence.

**Figure 2:**
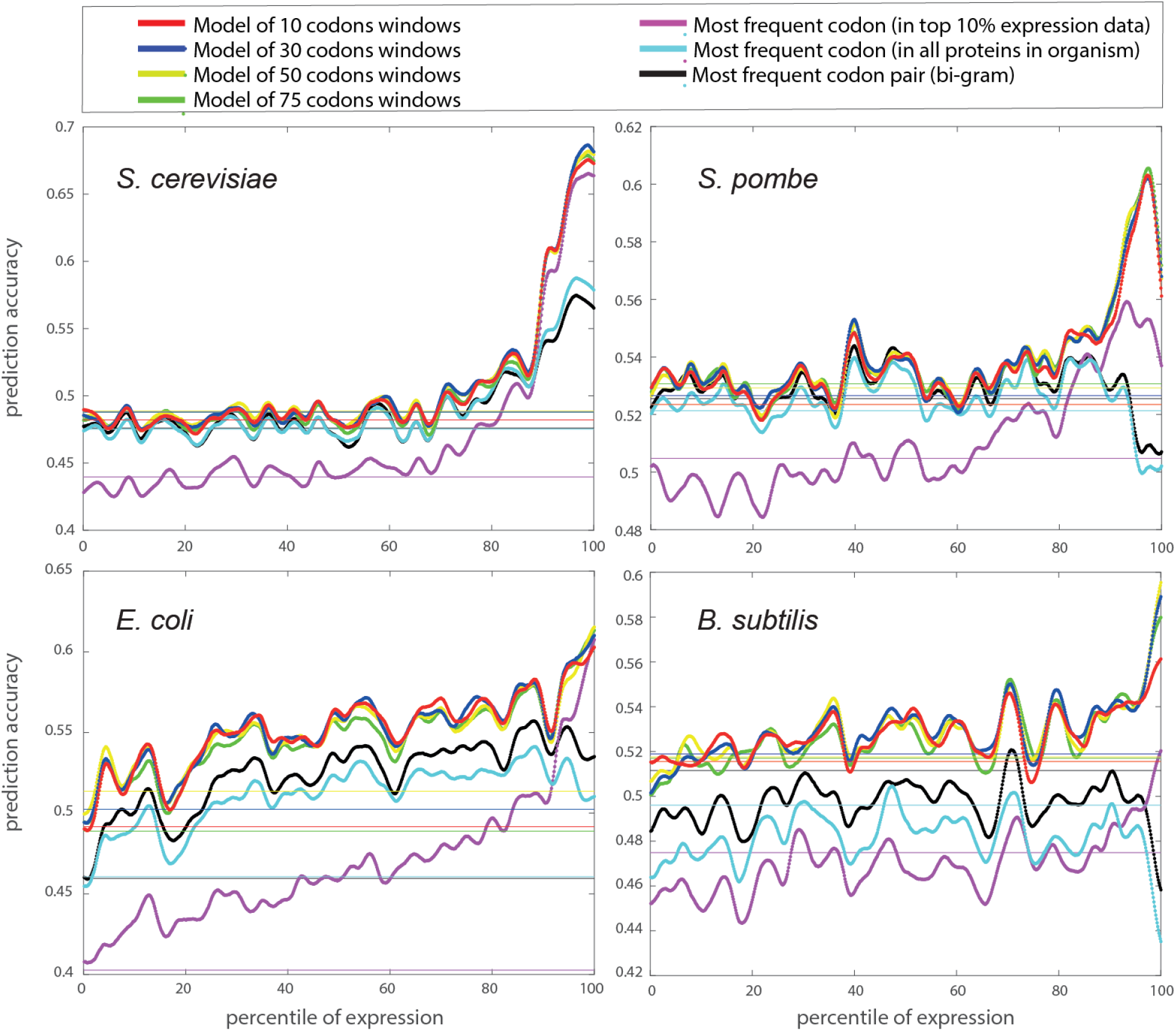
The codon prediction accuracies for the test set proteins with inference in masking mode show that mBART trained models predicts codons better than the frequency-based models; the model with a 30 codons window is generally the top-performer. Prediction accuracies for proteins in the test sets of *S. cerevisiae, S. pombe, E. coli*, and *B. subtilis* plotted versus percentile ranking of expression. The average accuracies for proteins for which expression has not been measured are shown as solid horizontal lines. Data were smoothed with a Gaussian filter and a window size of 50 proteins. The mBART masking models with 10, 30, 50, and 75 codon windows are shown in red, blue, yellow, and green, respectively. The frequency-based model accuracies calculated on all data are shown in cyan, frequency-based model accuracies calculated on the top 10% of proteins based on expression levels are shown in magenta, and the bigram frequency-based model in black. In all four organisms, predictions improve when considering proteins that are more highly expressed. The improvement in accuracy is most pronounced when considering the bacterial proteins. That the accuracy of our models is better than the frequency-based baselines demonstrates that there is an evolutionary pattern of long-range dependencies among codons, and it is sufficiently pronounced that the AI models considered here can learn it.

### (2) Masking-mode mBART predictions have better accuracy than the frequency-based baseline suggesting that long-range codon interaction patterns can be learned

Figure 2 shows the accuracies of codon predictions for the test-set proteins in masking mode for different models, as a function of expression level in the four organisms. The mBART models were pre-trained on the masking and mimicking tasks and then fine-tuned (FT) on the masking task. The models varied by window size, with windows of 10, 30, 50, and 75 codons evaluated. We fixed the mBART window size in each model so that we could reason about the scale of the learnable long-range interactions. The accuracies of the baseline frequency-based models, which are based on the most frequently used codon for each amino acid in each organism for all training-set proteins (cyan), in the 10% most highly expressed training-set proteins (magenta), and the frequency-based bigram model (black) were calculated for comparison. The mBART models are more accurate than the frequency-based models, suggesting that there are patterns that can be learned from the long-range relationships among codons.

We sort the proteins by expression level and show their ranked position along the *x*-axis; Figure S1 shows the same data with expression values along the *x*-axis. In both cases, the data is smoothed with a Gaussian kernel (50 proteins window), and the horizontal solid lines indicate the average accuracy for the proteins with no measured expression. Figures S2, S3 show the accuracy differences between our models and the frequency-based baselines. Tables S1 and 2 list the accuracy differences, the p-values, and the effect size (normalized Cohen d-values) comparing pairs of models. Table S1 compares different mBART models, showing that the 30 codons window performs best; Table 1 compares the 30 codons window to the frequency-based models. The statistical test considers the null hypothesis that the data in the differences (e.g., accuracy of prediction of the mBART model minus the accuracy of the frequency-based model) comes from a normal distribution with zero mean and unknown variance, using the paired-sample t-test. All p-values for comparisons with the baseline models are (far) smaller than 0.05 (Table 1), thus rejecting the null hypothesis, and suggesting the accuracies of the mBART predictions are indeed better than that of the frequency-based predictions.

**Table 1:** Significance of differences in accuracies between models. P-values and Normalize Cohen d values were determined using a paired-sample t-test on the test set proteins (grouped by organism) for the test decision for the null hypothesis that differences between models comes from a normal distribution with mean equal to zero and unknown variance. We see that in all cases, there is a difference, as evidenced by the absolute mean percent, the p-values that indicate that the difference is significant and the normalized Cohen d value indicating that the effect size is meaningful.

| Compared predictions | *S. cerevisiae* |  |  | *S. pombe* |  |  |
| --- | --- | --- | --- | --- | --- | --- |
|  | p-value | effect size |  | p-value | effect size |  |
|  |  | Absolute (<% diff>) | Normalized (Cohen d) |  | Absolute (<% diff>) | Normalized (Cohen d) |
| mBART (ws 30) vs. frequency-based (all) | <10<sup>-5</sup> | 1.5395 | 0.2047 | <10<sup>-5</sup> | 1.2272 | 0.2830 |
| mBART (ws 30) vs. frequency-based (bigram) | <10<sup>-5</sup> | 1.5332 | 0.1791 | <10<sup>-5</sup> | 0.7951 | 0.1602 |
| mBART (ws 30) vs. frequency-based (top 10%) | <10<sup>-5</sup> | 4.2307 | 0.9644 | <10<sup>-5</sup> | 2.7519 | 0.7327 |
|  | *E. coli* |  |  | *B. subtilis* |  |  |
|  | p-value | effect size |  | p-value | effect size |  |
|  |  | Absolute (<% diff>) | Normalized (Cohen d) |  | Absolute (<% diff>) | Normalized (Cohen d) |
| mBART (ws 30) vs. frequency-based (all) | <10<sup>-5</sup> | 4.0541 | 0.7363 | <10<sup>-5</sup> | 4.5887 | 1.1914 |
| mBART (ws 30) vs. frequency-based (bigram) | <10<sup>-5</sup> | 2.8260 | 0.4966 | <10<sup>-5</sup> | 3.0508 | 0.8132 |
| mBART (ws 30) vs. frequency-based (top 10%) | <10<sup>-5</sup> | 8.4185 | 1.4942 | <10<sup>-5</sup> | 5.8368 | 1.4337 |

Figure 3 compares the perplexity of different models on the test-set proteins, as a function of the expression level rank of the proteins (with a Gaussian kernel smoothing of 50 proteins). Perplexity is a commonly-used measure in AI to assess model predictions and is the computed exponentiated average of the cross-entropy loss, implying that better models have lower perplexity. The intuition behind perplexity is modelling the “surprise” of a model with respect to the naturally-occurring codon sequences in the test sets, in terms of the probability the models assign these sequences. Namely, it complements the accuracy measure, which only rewards cases where the model assigns the highest probability to the correct codon.

**Figure 3:**
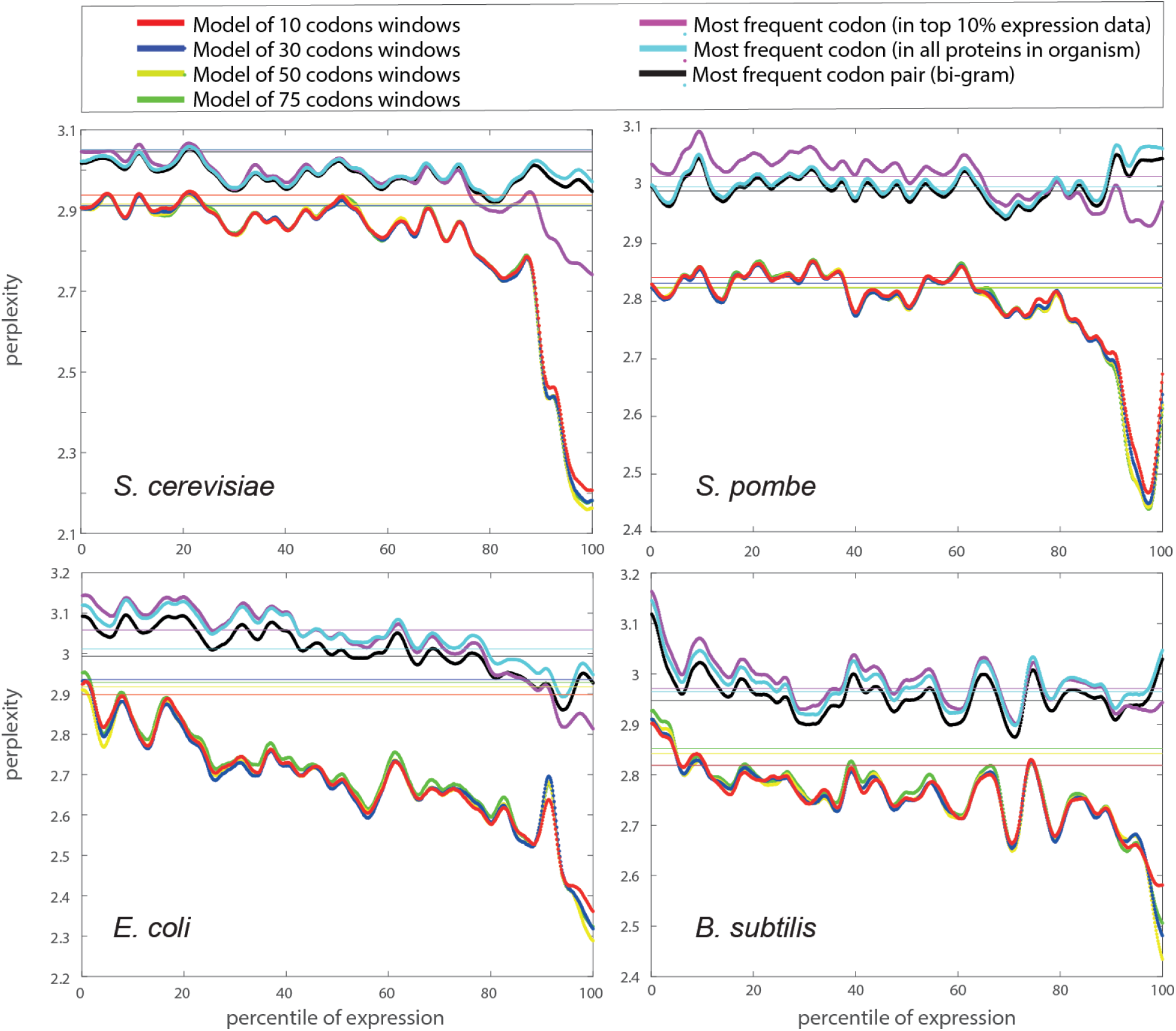
Calculated perplexities for masking-mode mBART predictions are lower than those of the frequency-based models. Perplexity, which is the computed exponentiated average of the cross-entropy loss, plotted versus percentile ranking of expression. The average perplexities for proteins with no measured expression levels are shown by solid horizontal lines. The data were smoothed with a Gaussian filter, and a window of 50 proteins. These graphs complement Figure 2 showing the accuracies: The mBART models perform better (lower perplexity) than the frequency-based models, the model with a window size of 30 codons is a top performer, and the perplexities are lower when considering proteins that are more highly expressed. This provides further support to that there is an evolutionary pattern of long-range dependencies among codons, and it is sufficiently pronounced that the AI models considered here can learn it.

Figures 2, 3, S1-S3 and Tables S1,2 show that the mBART models predict more accurately than the frequency-based models, demonstrating there are patterns that can be learned from the long-range relationships among codons. Both for the frequency-based models and the mBART models it is easier to predict the codons of more highly expressed proteins. In *S. cerevisiae*, the advantage of mBART is the smallest and is in the highly expressed proteins. In *S. pombe*, the best mBART models are better than the frequency-based models, and for the bacteria, the mBART models offer the largest improvement in prediction accuracy and perplexity with respect to the frequency-based models, across all levels of protein expression. Furthermore, considering both the accuracy and the perplexity shows that the AI-based models can learn patterns of codon usage even better when relying on a window that is even longer than 10 codons.

We also evaluated the accuracies of the mBART and the frequency-based models as a function of the protein length ranking (Figures 4, S4-6). There is not a consistent relationship between the accuracy of the best model and protein length. In *S. cerevisiae, E. coli*, and *B. subtilis*, longer proteins are more accurately predicted; in *S. pombe* this is not the case. The accuracy advantage of the mBART models is more pronounced in the shorter proteins of the eukaryotic organisms and the longer proteins of the bacterial organisms (Figure S5-6).

**Figure 4:**
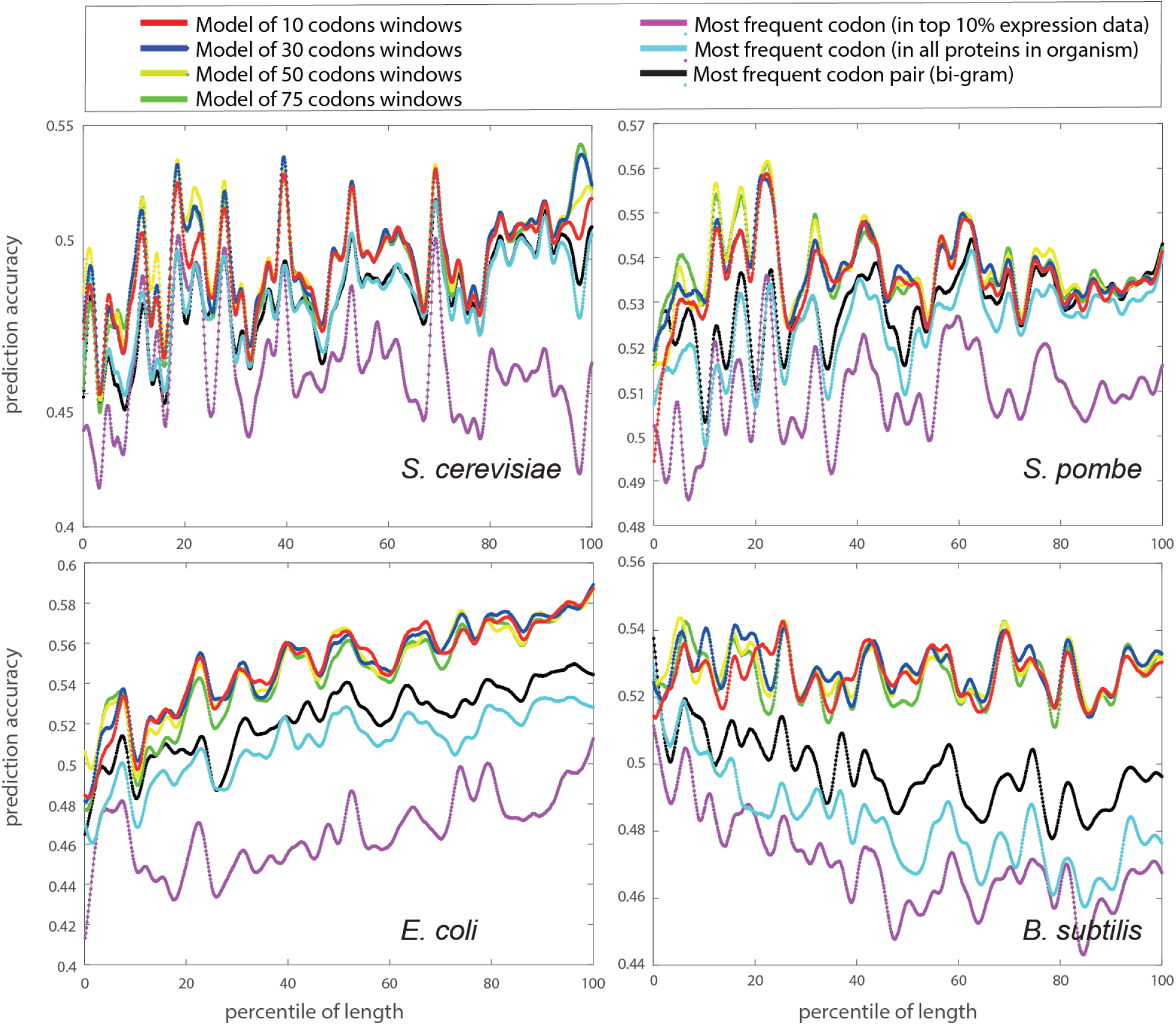
Codon prediction accuracies for the test set proteins with inference in masking mode do not have a clear dependency on length. Prediction accuracies versus length percentile ranking, with the data smoothed with a Gaussian filter (window size of 50 proteins). There is no simple connection between the accuracies of the models (any of them) and the lengths of the proteins: in *S. cerevisiae* and *E. coli* the mBART models are more accurate for longer proteins, in *B. subtilis*, the models are more accurate for the shorter proteins, and in *S. pombe*, there is no clear dependency between the two.

To determine whether similarities between the training and test sets artificially boost prediction accuracies, we identified similarities at the amino-acid level that remained after we used CD-HIT clustering for the training set/test set split. To do this, we BLAST aligned (with an E-value threshold of 10^-2^) each test protein to the training set. Then, we calculate its average percent identity to the training set by identifying for each residue the aligned segment with the highest percent identity and averaging these values over the protein residues. This analysis showed that 31% of the test set proteins have no sequence identity to proteins in the training set (28.1%, 28.1%, 30.3%, and 37.2% for *S. cerevisiae*, *S. pombe*, *E. coli*, and *B. subtilis* respectively; Figure S8). The accuracies of the predictions of the mBART and frequency-based models as a function of the average identity were then calculated (Figure S9). The accuracy gap between the mBART and frequency-based models was similar regardless of whether or not the test set protein had close neighbors in the training set, suggesting that the performance of the mBART models is not artificially boosted by these similarities.

### (2) Mimicking-mode mBART predictions are only marginally better than masking-mode predictions

To identify orthologous segments that can be used in mimicking-mode predictions, we used BLAST (with an E-value cutoff of 10^-2^). Figure S10 shows histograms quantifying the similarities among the orthologous segments in the test-set using percent identity and log_10_(E-values). The average percent identities among the orthologous segments in the test set are 32%, 33%, 30%, and 29% for *S. cerevisiae*, *S. pombe*, *E. coli*, and *B. subtilis*, respectively. During training, the data is of aligned proteins that are both from the training set, and in testing, orthologous proteins that were both in the test set were evaluated.

We evaluated different mBART models in masking-mode inference and mimicking-mode inference, on the test set of orthologous segments. First, we consider the same models (FT on the masking-task with windows of 30 and 50 residues) and the same masking-mode inference as described above, only on this different test set. We further fine-tuned these two models on the masking and mimicking task, and evaluated these refined models in masking-mode inference and in mimicking-mode inference. Finally, we also calculated the frequency-based baseline (similarly to the what is described above, only on this test set), and a naïve frequency-mimicking model where we mimic the codon with the same frequency rank as in the orthologous protein (see Methods for details). Figures 5 shows the accuracies of the predictions as a function of the sequence identity to the orthologue; Figures S11 shows these accuracies with respect to the frequency-based baseline, and Figure S12 shows the same data, sorted by accuracy along the *x*-axis.

**Figure 5:**
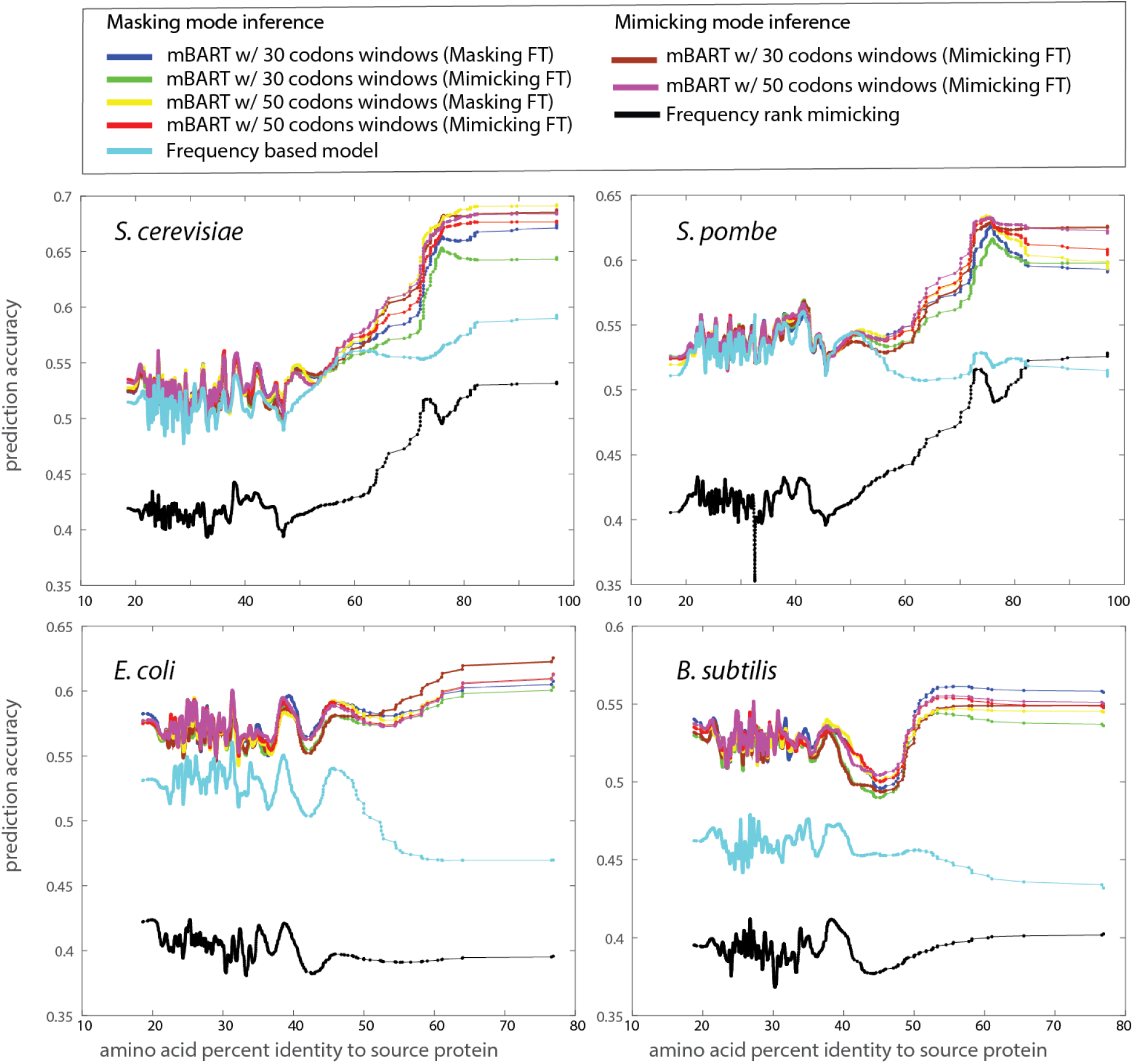
mBART mimicking-mode inference accuracy is on par with the masking mode inference. Prediction accuracies versus amino acid percent identify to the orthologous protein. We use two inference modes: masking and mimicking and different models. In masking-mode, models that were fine-tuned on the masking task with window sizes of 30 and 50 are shown in blue and yellow respectively. These are the same models and inference mode shown in Figure 2, only here they are evaluated on the protein segments found in the alignment dataset. We also further fine-tuned these models on the masking and mimicking task, and the accuracies of the predictions of these FT models in masking-mode inference are shown in green and red, respectively. Alternatively, we used these same models in mimicking-mode inference and the accuracies of these predictions are shown in maroon and magenta. For comparison, we show the frequency-based model on this dataset in cyan, and the prediction accuracy of a frequency-based mimicking model in black. Data were smoothed with a Gaussian filter (window size of 50 proteins). The signal of codons in the orthologous proteins is not sufficiently strong that our AI models can exploit it to improve their predictions, and the performance of the mimicking-mode inference (in maroon/magenta) is on par with that of the masking mode inference (in green and red), with a slight advantage for proteins with very close orthologues in bacterial organisms.

The mBART model’s mimicking-mode predictions have accuracies that are very similar to the masking-mode predictions. The mimicking-mode resulted in a modest accuracy boost in the two eukaryotic species and for orthologs that have segments with high amino-acid sequence identity. Prediction accuracy may be higher due to the codons used in a particular segment are easier to predict, namely the most frequently used ones. In the masking-mode frequency-based predictions, ortholog codons are not used, and hence percent identity is only meaningful in terms of characterizing the protein as a conserved one. In the eukaryotic species, the accuracies of both the naïve frequency-based mimicking and the mBART models became higher than the frequency-based baseline as the percent identity of the orthologs increased. This implies that there is a signal in the codons of the orthologs, especially when the percent identity is high, that improves prediction accuracy. However, the masking-based predictions have almost identical accuracies, so this extra information does not further improve the mBART predictors. It should be noted that even when there is a signal, the information from the orthologs could be detrimental. For example, although the naïve frequency-based mimicking improved due to this signal, it was still less accurate than the frequency-based masking model. Our mimicking-based predictions have similar, and in some cases, improved accuracy compared to the naïve frequency-based mimicking model.

Table S2 shows the accuracy differences, p-values, and effect sizes (normalized Cohen d values) comparing pairs of models. The comparison of the mBART model in mimicking mode and the frequency-based mimicking model for all four organisms shows that the difference is significant (e.g., all p-values are less than 10^-5^). In contrast, the comparison of the mBART model in masking mode versus in mimicking mode shows their differences are not significant.

### (3) mBART outperforms the frequency-based baseline more significantly for certain types of proteins

To determine whether prediction accuracies depend on molecular functions or type of biological process, we use the GO-XL slim classifications of *S. cerevisiae* and *E. coli* proteins and considered cases for which there were at least 10 proteins with that annotation in the test sets of these organisms. For each annotation, placed along the *x*-axis, and the data is shown as box plots and scatter data. We used the Mann-Whitney rank sum test to determine if the accuracy difference values for the proteins with a specific GO annotation are likely from the same population of values as the rest of the organism’s test-set proteins (Table S3).

For most GO-annotation categories, the accuracy boost of the mBART model is not different from the background of all test-set proteins in either organism, but there are some that stand out (Figures 6 and S13-S16). In *S. cerevisiae*, the molecular function catagories in which the mBART model predicts even better than the frequency-based baseline are: (1) ‘structural constituent of ribosome’, (2) nucleic acid binding function ‘rRNA binding’, ‘RNA binding’, and ‘DNA binding’, and (3) molecular function that falls under the broad category of catalytic activity and specifically ‘transferase activity’, ‘nucleotidyltransferase activity’, ‘nuclease activity’, and ‘peptidase activity’. For *E. coli,* these are: (1) ‘structural molecular activity’, (2) ‘RNA binding’, and (3) both the broader ‘catalytic activity’ and more specifically ‘catalytic activity acting on DNA’. Considering biological processes, in *S. cerevisiae* these are: (1) ‘ribosomal large subunit biogenesis’, (2) the broad category of translation and specifically ‘cytoplasmic translation’ and ‘regulation of translation’, (3) the broad category of nucleic acid metabolic process and specifically ‘DNA recombination’ and ‘rRNA processing’, and (4) ‘transposition’. In *E. coli:* (1) ‘ribosome biogenesis’, a parent category of ‘ribosomal large subunit biogenesis’ identified in *S. cerevisiae*, (2) ‘cytoplasmic translation’, a process also identified in *S. cerevisiae*, (3) and ‘protein containing complex assembly’.

**Figure 6:**
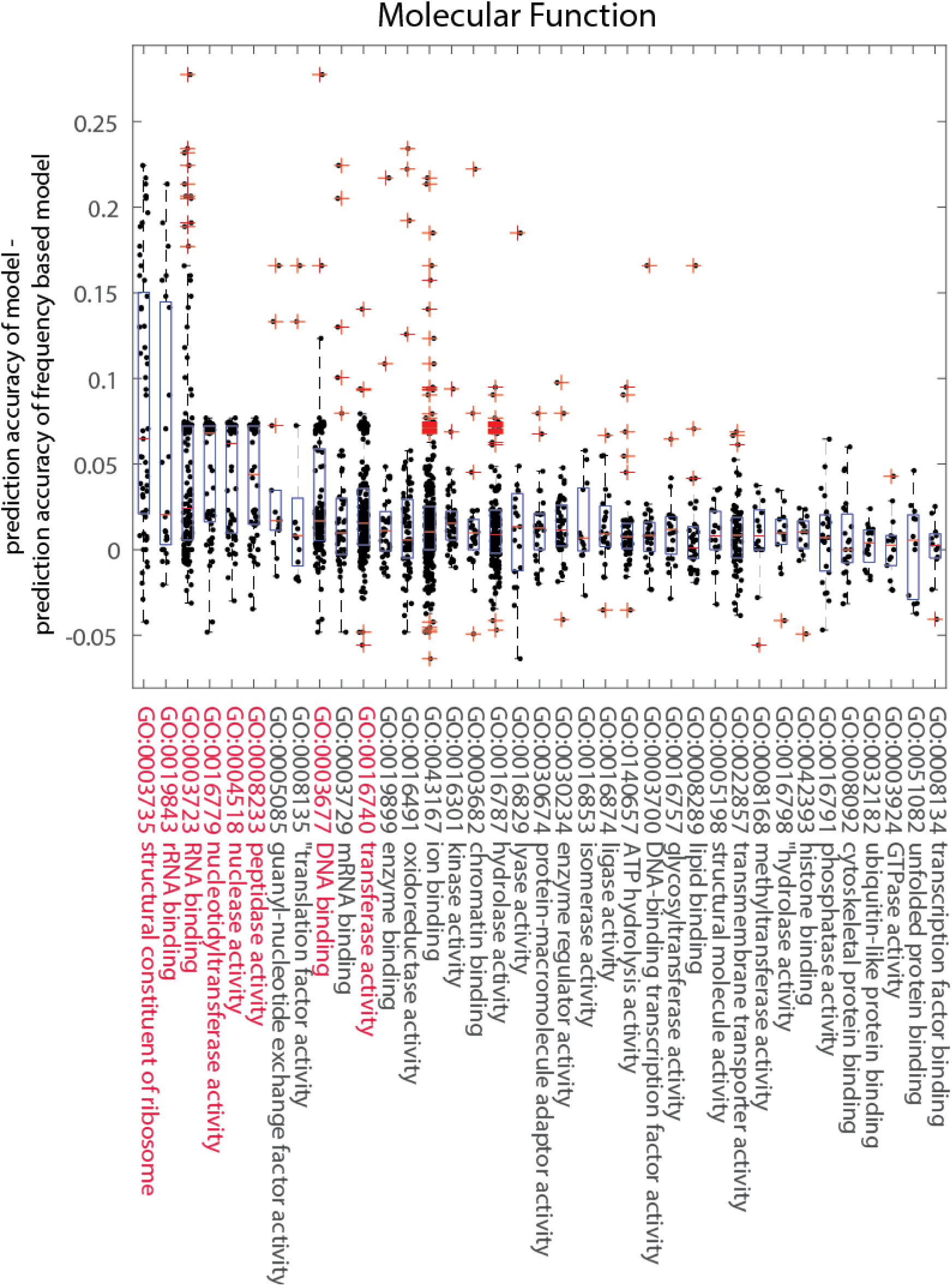
Prediction accuracy is high for proteins in certain GO functional groups. Differences between masking-task-FT mBART model (30-codon window) predictions and the frequency-based baseline for *S. cerevisiae* test-set proteins grouped by GO molecular function terms. Terms were sorted in descending order by the mean average difference. The terms for which the p-value is <0.05 in a Mann-Whitney rank sum test for the functionally grouped set versus the rest of the test-set proteins are highlighted in red. We see that the mBART model outshines the naïve approach more significantly for proteins with the function ‘structural constituent of ribosome’, nucleic acid binding function, and sub-functions of catalytic activity.

## Discussion

We have employed contemporary tools from AI to predict codons in genes from the eukaryotes *S. cerevisiae* and *S. pombe* and the bacteria *E. coli* and *B. subtilis*, given the protein amino acid sequences. Our approach is data-driven: mBART models were trained on data from each organism and are evaluated on a separate and diverse test set. Because the difficulty of codon prediction varies, evaluation on a large and diverse test set rather than on only a few proteins as has been done previously [26] results in a more comprehensive test of predictive power and allows us to study the accuracy of the models as a function of protein expression, length, conservation, and functionality. The lower bound for the accuracy of the model are the frequency-based models, and we expect the effective upper to be significantly lower than 100%, because not all positions are under a strong evolutionary pressure [49]. Given this, the improvement reported here relative to the frequency-based model is significant. The datasets we used for training/testing and our mBART models are publicly available, both as source code and trained models and through a web-based user-friendly interface (https://www.aa2codons.info/).

Learning the statistical patterns of codon usage in an organism is challenging because the amount of available data is limited by the number of proteins in the organism. Previous studies highlighted associations between the frequencies of neighboring codons and attributed these to their effect on ribosomal pausing, frameshifting, and other gene expression steps [30, 51, 52]. Here, we used mBART, a transformer-based architecture [31], to learn correlations in codon use in residues that are 10 to 75 codons apart to improve predictions. Codon usage patterns inferred by our models are therefore related not only to translation but also to gene expression steps including transcription, splicing, methylation, RNA processing, mRNA stability, and genomic stability [7]. Each of our models relies on a fixed window size, allowing us to reason about the distance among codons in which our models can still improve, presumably by learning patterns among codons that are separated this far apart. That the model with a 30-codon window was a better performer than models with longer window sizes may be due to statistical aspects such as the amount of data given the number of parameters in the model. However, this window size may be optimal because it is closest to the length of the ribosomal exit tunnel [53].

That our trained models can better learn the codon patterns of a subset of proteins, suggests that their codon usage is more constrained, and this may be due to the encodings of these proteins being under more pronounced evolutionary selection. Here, we study how the accuracy and perplexity of our trained models vary as a function of the expression-levels, lengths of the proteins, conservation, and GO annotations. It was suggested that selection for codon usage increases with the expression levels of a gene [25, 54]. One reason for this is that we expect a silent mutation to have a greater effect on organism fitness in highly expressed genes [25]. Indeed, the codons of high-expression proteins in all four organisms, are more accurately predicted both by the naïve frequency-based models and our models, and with lower perplexity. This suggests that high-expression proteins are not only under increased evolutionary selection for simple codon patterns but also for complex-long range codon usage patterns. We also observed that in most organisms (excluding *S. pombe*) the gap between the performances of our models and the frequency-based approach is greater for long proteins than for the short ones. It is possible that complex signals related to co-translational folding regulation, needed for the tighter control of in multi-domain proteins [55, 56], are “encoded” by codon patterns at distant positions and thus better detected by our models. It is also possible that our models detect gene expression codes beyond those important for co-translational folding that are encoded in longer genes; these could be codes related to binding sites of transcription factors or RNA binding proteins [7].

Our models performed better for genes with conserved orthologs than for those without. The latter are either new or have undergone very rapid evolution. This suggests that evolution first shapes the amino acid sequence and later the complex codon usage patterns. It may also suggest that older genes tend to include more complex gene expression codes than newer ones, possibly as a result of selection for tighter regulation, and it is these complex codes that are detected and exploited by our models.

The gap between the performances of our models and the frequency-based baselines varied among different GO functional groups. This result may be related to factors mentioned above and that seem to introduce complex codes into the coding sequence: expression levels, gene age and conservation, and gene length. For example, genes related to the translation process (e.g., those encoding ribosomal proteins), that were better predicted by our models, are known to be highly expressed and old. DNA and RNA binding proteins are also known to include ancient domains, and this may explain the better performance of our models in predicting their codons [57]. Our GO-based analysis suggests that our models can be useful for studying and annotating novel genes given that predictive performance is associated with functionality.

Finally, our models predict better than the frequency-based approach more significantly in bacteria than in the yeast species evaluated. This may be due to the larger effective population size in bacteria [58]. A larger population should induce stronger selection pressure on codon usage, which in turn will result in complex-long-range signals that can be learned by our models but not by the frequency-based models. Also, the mean number of ribosomes per mRNA in bacteria is greater than in eukaryotes making traffic jams more common, and possibly triggering stronger selection for complex codon usage signals [59–62]. Moreover, horizontal gene transfer occurs in bacteria [63], which may accelerate the evolutionary rate.

We trained our mBART models to mimic, i.e., to predict the codons of a protein given the codon encodings of an ortholog. An accurate mimicking tool is useful for predicting codons that are optimal in a non-native host. To design a mimicking tool for protein codons, the first step is to characterize patterns of codon usage in orthologous proteins. Then, the learned insights (or AI-models) can be used, given the codon encoding of a protein in a source organism, to predict a codon encoding for a similar protein in a target organism. The final, sometimes overlooked step, is to evaluate the predicted codon encodings in the target organism.

Previous studies used “hand-crafted” measures to characterize patterns of codon usage in orthologous proteins: Pechman and Frydman devised a translational efficiency scale and applied it to yeast species to show evolutionary conservation of codon optimality in eukaryotes [32]. Jacobs and Shakhnovich measured local rare-codon enrichment and studied its conservation across multiple-sequence alignments [33]. Chaney et al. used the MinMax measure to identify clusters of rare codons and showed that they are conserved among homologous proteins across eukaryotic and bacteria species [64]. These measures are only a few of the many available codon bias indices [25]. In contrast to these hand-crafted features, our mBART models learn patterns from the data, both indirectly by training a single model on the masking task for multiple organisms and directly by training on the mimicking task from alignment data. As in all contemporary AI-based models, the patterns learned help improve accuracy during inference, even though we do not have an explicit description of what the models learned.

Evaluating mimicking tools is challenging because there is a discrepancy between their desired use and the data on which we can evaluate them. Their use is to predict the optimal codons for expression of a protein in a non-native host. Evaluation can be carried out on evolutionary data: the codon encodings in two organisms of merely similar, namely non-identical, proteins. This meaningful distinction has two important consequences. First, because evolutionary data does not include pairs of identical protein sequences in two organisms as in our desired use, current tools, like CHARMING [36] and CodonWizard [35], do not evaluate their measures, or even predict codons, based on the codon encodings of a similar, yet non-identical, protein ortholog. Second, it is unclear what is a correct threshold for approximating this desired use, in terms of the amino-acid sequence identity of the ortholog. We observed that mimicking predictions are not consistently better than the masking predictions. This may be because we evaluated our mimicking models on orthologs that are too remote. Alternatively, it may be that it is not easier to learn to mimic than to predict the codon encoding from the amino acid sequence. Nonetheless, accuracies of our mimicking-mode predictions improved as the amino acid sequence identity increased, approximating the desired use more closely. Thus, we believe that our mimicking-based predictions will be useful for optimizing expression of proteins in a non-native host, and we provide the code and a web-interface for this task.

In summary, we used AI to study codon usage bias. Our approach can be used to design the coding sequence of heterologous genes and also to study where there are learnable patterns of codon encodings. We believe that that future research on codon usage will involve extensive application of AI-based approaches, like the one presented here.

## Acknowledgements

We thank Prof. Rita Osadchy of University of Haifa and Michael Peeri, Lorna Bakhit, and Prof. Nir Ben-Tal of Tel-Aviv University for insightful discussions. SBE and TT were supported by the Safra Center for Bioinformatics at Tel-Aviv University, and TS and RK were supported by the Data Science Research Center at the University of Haifa.

## Methods

<u>Dataset:</u> The NCBI coding sequences of four organisms, *S. cerevisiae*, *S. pombe*, *E. coli*, and *B. subtilis*, were divided to three disjoint sets, for training, validation, and testing. We first clustered their translated amino acid sequences with CD-HIT (flags -d 10000 -c 0.7 -bak 1). Clusters were not split between two sets (e.g., training and test); the resulting sets are available online (https://www.aa2codons.info/). Within each of the three sets (i.e., training, validation, test), we used BLAST (with E-value threshold 0.01) to identify similar sequences. Finally, we associated with each protein, if available, measured expression levels [65], and its category in the 6-class classification: [0-25%, 25-50%, 50-75%, 75-90%, 90-100%, no measured expression].

<u>Models:</u> We predict the codons using three frequency-based models: (1) for each organism, for each amino acid, the most frequently used codon in the test set (2) for each organism, for each amino acid, the most frequently used codon in its top 10% highly-expressed proteins (3) a bi-gram table calculated for each organism, for each amino acid, the most frequently used codon conditioned on its preceding codon. We also considered the bi-gram model with a frequency table that is conditioned on the preceding amino-acid, and found that its predictions are less accurate.

We trained and share mBART models with fixed window sizes (10, 30, 50, and 75) using the Huggingface infrastructure [66]. mBart has an Encoder-Decoder architecture that receives two inputs: one for the encoder and one for the decoder; the decoder is auto regressive. We tested models with this architecture that vary in their specific configuration: the number of attention layers (3, 6, 12), the number of attention heads (4, 8, 16), the dimension of the feed forward network (64, 256, 1024) and the d_model (32, 64, 128). Based on this hyper-parameter search, we chose to train models with six attention layers and eight attention heads for both encoder and decoder. The auto-regressive nature of the decoder means that it predicts probabilities for the next codon, given the encoder inputs and all the previous predictions. To reduce training time, we used teacher-forcing, where the decoder predicts probabilities for the next codon given the input to the encoder and the correct previous predictions (as opposed to its own predictions which would be given in non-teacher-forcing mode). Critically, during inference, the codon sequence is not given, and a fixed-length window of prediction with the same size of the input is generated auto-regressively choosing the highest probability codon restricted to the amino-acid sequence we want to generate.

<u>Training:</u> Our training protocol had three steps: pretraining, masking-mode fine-tuning, and mimicking-mode fine-tuning. There are hyper-parameters that govern the training procedure, and we searched amongst those as well: we considered 1e-4, 1e-5 learning rate, different masking percent (10%, 30%, 50%, 70%), different batch sizes (8, 32, 128, 512), different numbers of warmup steps (500, 5000). Based on this search, we chose the following training procedure: Each training step had a batch size of 32, 15000 warmup steps, 1e-4 learning rate, 15% label smoothing, and early stopping on the cross-entropy loss of the validation set. In pretraining, we randomly chose 30% of the input codons and masked each with a token of its encoded amino acid. The training set windows were randomly chosen (both the sequence and the position within the sequence) on-the-fly during the batch creation. To avoid noisy comparisons between validation steps that are used for early stopping, we pre-calculated and fixed the validation set windows. In mimicking-mode training, we only masked the codons of the aligned target and noted that these may differ from the amino acids of the orthologous sequence. Figure S7 shows as an example of the cross-entropy loss on the training set (train/eval) for the pre-trained and fine-tuned models with a window size of 30 codons.

<u>Inference:</u> Testing during inference was on coding sequences of a wide range of lengths, yet our mBART models train (only) on fixed-sized windows. Naively applying the models trained on fixed-sized windows to longer sequences led to poor performance. Thus, we used our models only to infer predictions in windows of the size on which they were trained and derived a prediction for the full-length sequence from these partial predictions. Given a sequence and a model trained on a fixed window size, we predicted for all (sliding) windows of that size in the full sequence (using the ‘generate’ function). Then, for each position in the full-length protein, we averaged the logits predicted in that position in the sliding windows in which it appears and normalized these averages with log softmax. The final prediction was the highest probability codon among those that encoded the amino acid in that position.

<u>Frequency-rank mimicking:</u> For each position in the target sequence for which we wanted to predict the codon, we had a codon in the aligned orthologous segment. If the amino acid in the target sequence was aligned to a gap, then we predicted the most frequently used codon for that amino acid. If it was aligned to an identical amino acid, then we mimicked the rank frequency of the codon in the aligned position. For example, if the codon was the most frequently used codon in the other organism, we predicted the most frequently used codon in the target organism. Because it is the same amino acid, the number of possible codons is the same, and rank mimicking is straight-forward. Similarly, if the aligned amino acid in the other organism has the same number of codons encoding it as the target amino acid, rank mimicking is straight-forward. If the aligned amino acid has a different number of codons, we selected the codon with the highest-rank that was lower than or equal to that of the frequency in the mimicked codon.

<u>Evaluation metrics</u>: The prediction accuracy was calculated as the fraction of amino acids along the protein sequence that were correctly predicted based on the true, evolutionary selected, codon sequence of each protein. A perfect prediction has an accuracy of 1, and a prediction that failed in all positions has an accuracy of 0. We also measured the perplexity of a model given the true, evolutionary selected, codon sequence: Perplexity was computed by averaging the negative log(probabilities) assigned by the predictor to the true codon, and then exponentiating this average. In the case of the frequency-based baseline, the probability we assign each codon is its likelihood, and this is equal to exponentiating the average negative log likelihood.

<u>GO slim annotations for *E. coli* and *S. cerevisiae*</u>: GO slim annotations were downloaded from Quick GO (https://www.ebi.ac.uk/QuickGO/) on November 2023. GO terms were merged according to the GO slim data (all annotation related to one GO slim were merged). We used GO slim generic for *E. coli* and GO slim yeast for *S. cerevisiae.* Gene names were converted to unique names using UniPort ID mapping (https://www.uniprot.org/id-mapping). We removed from the analysis GO slim terms that contained less than 10 genes.

<u>Data availability</u>: Our models and datasets are available through the huggingface hub: models: https://huggingface.co/siditom, datasets: https://huggingface.co/datasets/siditom/SCPECBS3, tokenizer: https://huggingface.co/siditom/tokenizer-codon_optimization-refined_expr. A Google colab script for model inference (along with documentation) is also available: https://colab.research.google.com/drive/1ocbWMWcTHgGSQvgPtRuvY2CBxIVSHUxX?usp=sharing.

## Supplementary Material

**Figure S1:**
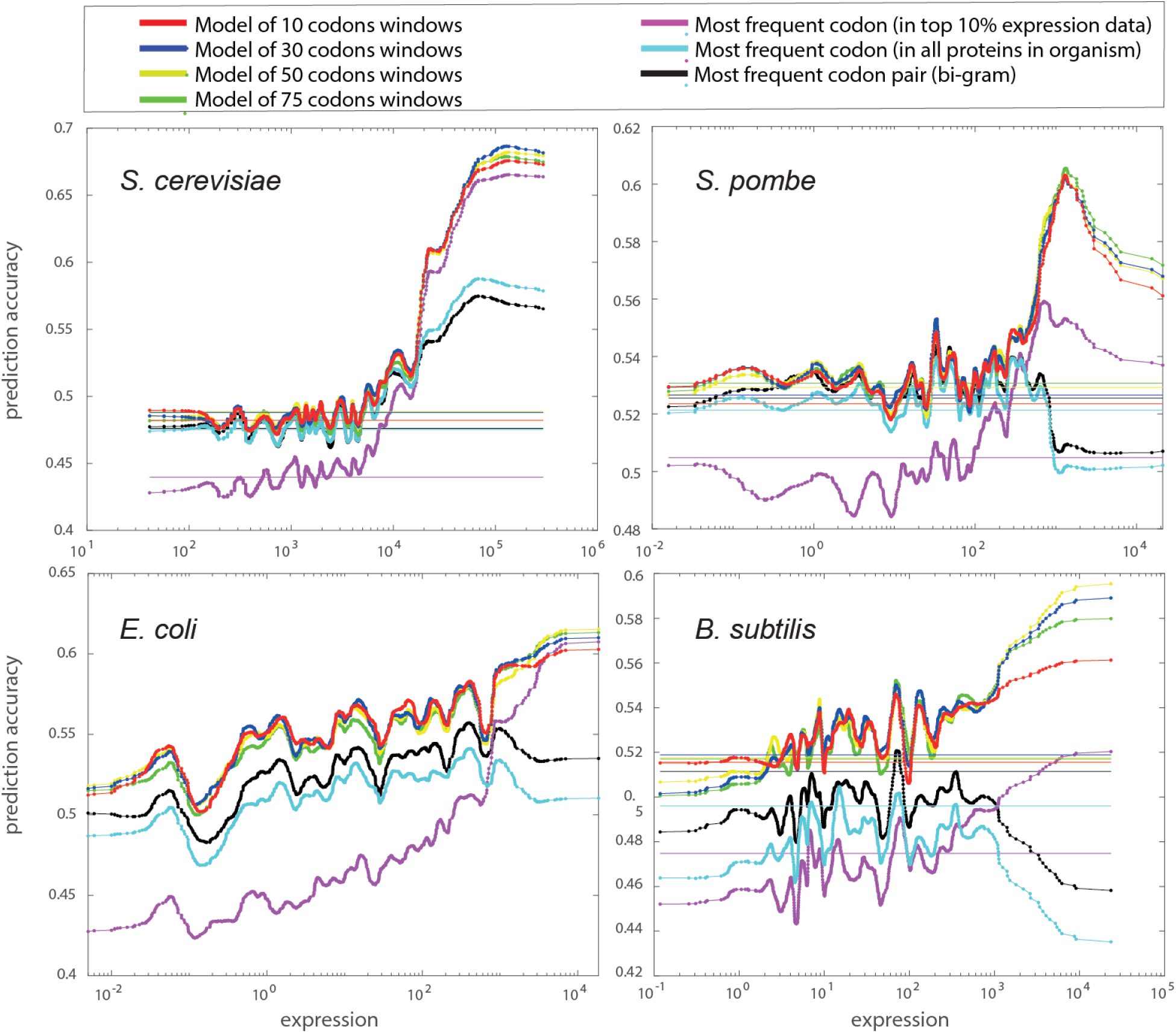
The codon prediction accuracies for the test set proteins with inference in masking mode versus log expression levels also show the model with a 30 codons window is the top-performer. Prediction accuracies are plotted versus expression level values in log scale. These are the same data as shown in Figure 2. The average accuracies for proteins for which expression has not been measured are shown as solid horizontal lines. The data were smoothed with a Gaussian filter (window size of 50 proteins). The mBART masking models with 10, 30, 50, and 75 codon windows are shown in red, blue, yellow, and green, respectively. The frequency-based model accuracies calculated on all data are shown in cyan, frequency-based model accuracies calculated on the top 10% of proteins based on expression levels are shown in magenta, and the frequency-based bigram model in black. In all four organisms, predictions improve when considering proteins that are more highly expressed. The improvement in accuracy is most pronounced when considering the bacterial proteins. That the accuracy of our models is better suggests that there is an evolutionary pattern of long-range dependencies among codons, and it is sufficiently pronounced that the AI models considered here can learn it.

**Figure S2:**
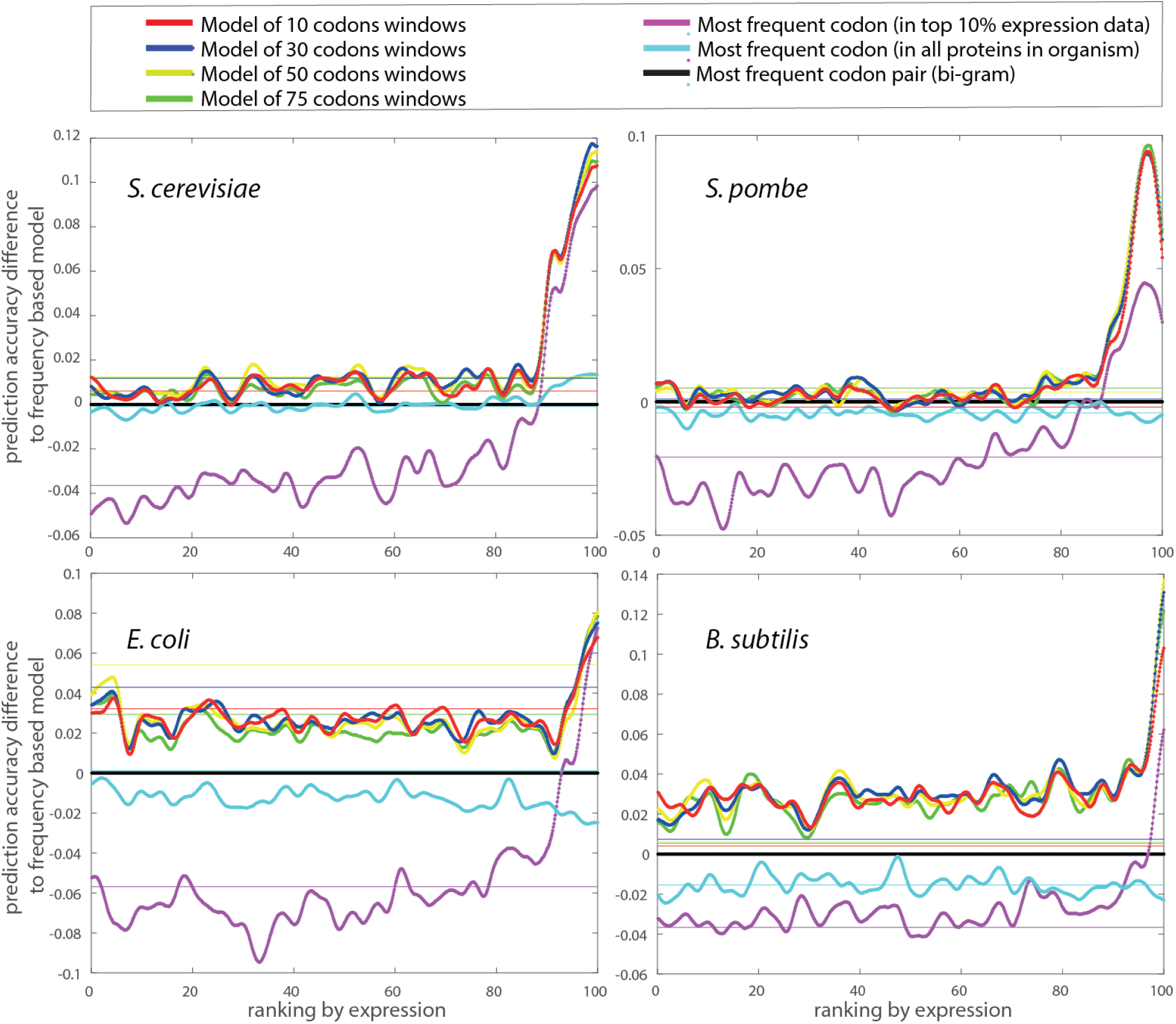
Differences in prediction accuracies from frequency-based bigram model. The differences in prediction accuracies between the mBART models and the frequency-based bigram model are plotted versus expression level rankings. These are the same data as in Figure 2. The frequency-based bigram model accuracies (black) are shown as a fixed value of 0. In this figure as well, we see that in all four organisms, the mBART models outperform the frequency-based baseline, and that the advantage is greater in the bacterial organisms.

**Figure S3:**
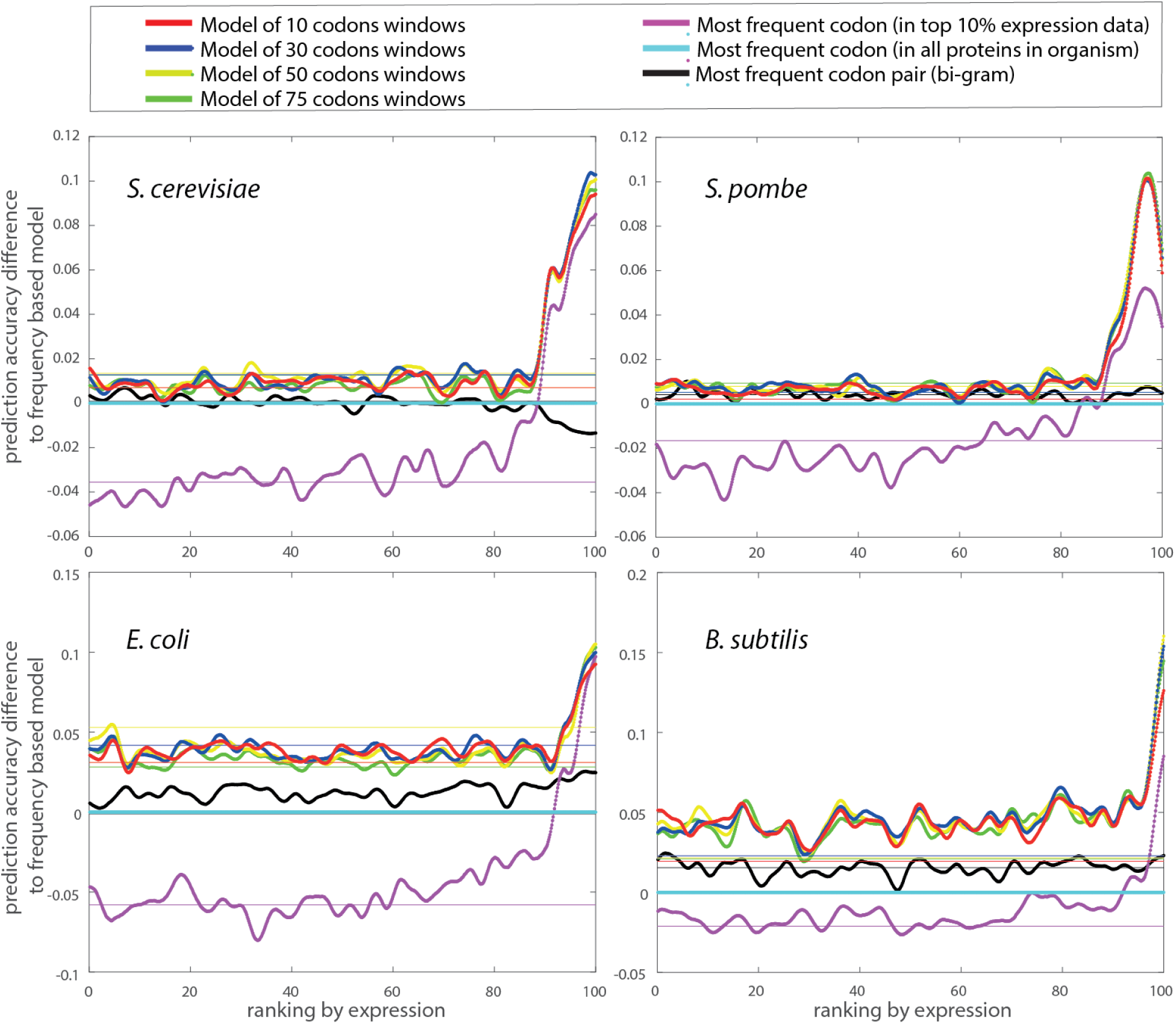
Differences in prediction accuracies from frequency-based model for all proteins versus ranking by expression level. The differences in prediction accuracies between the mBART models and the frequency-based model calculated on all proteins are plotted versus expression level rankings. These are the same data as in Figure 2, only the *y*-axis plots the differences in accuracies. Thus, the cyan frequency-based baseline is shown as a fixed value of 0. In this figure as well, we see that in all four organisms, the mBART models outperform the frequency-based baseline, and that the advantage is greater in the bacterial organisms.

**Figure S4:**
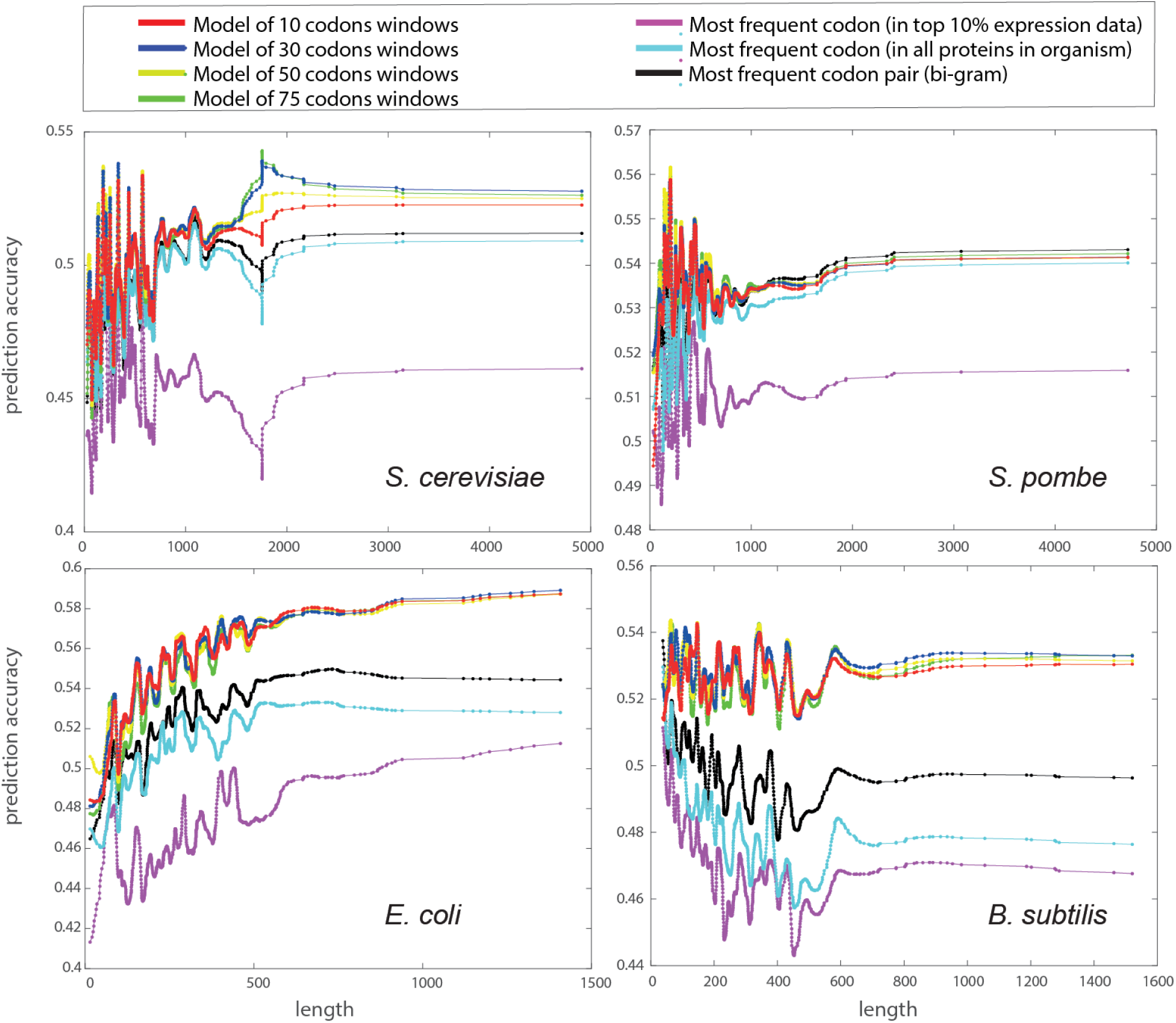
Codon prediction accuracies for the test set proteins with inference in masking mode with the proteins sorted by their length. The prediction accuracies between the mBART models and the frequency-based model plotted versus protein length. These are the same data as in Figure 4, but with actual protein lengths. Here too, we see that there is no simple connection between the accuracies of the models (any of them) and the lengths of the proteins: in *S. cerevisiae* and *E. coli* the mBART models are more accurate for longer proteins, in *B. subtilis*, the models are more accurate for the shorter proteins, and in *S. pombe*, there is no clear dependency between the two.

**Figure S5:**
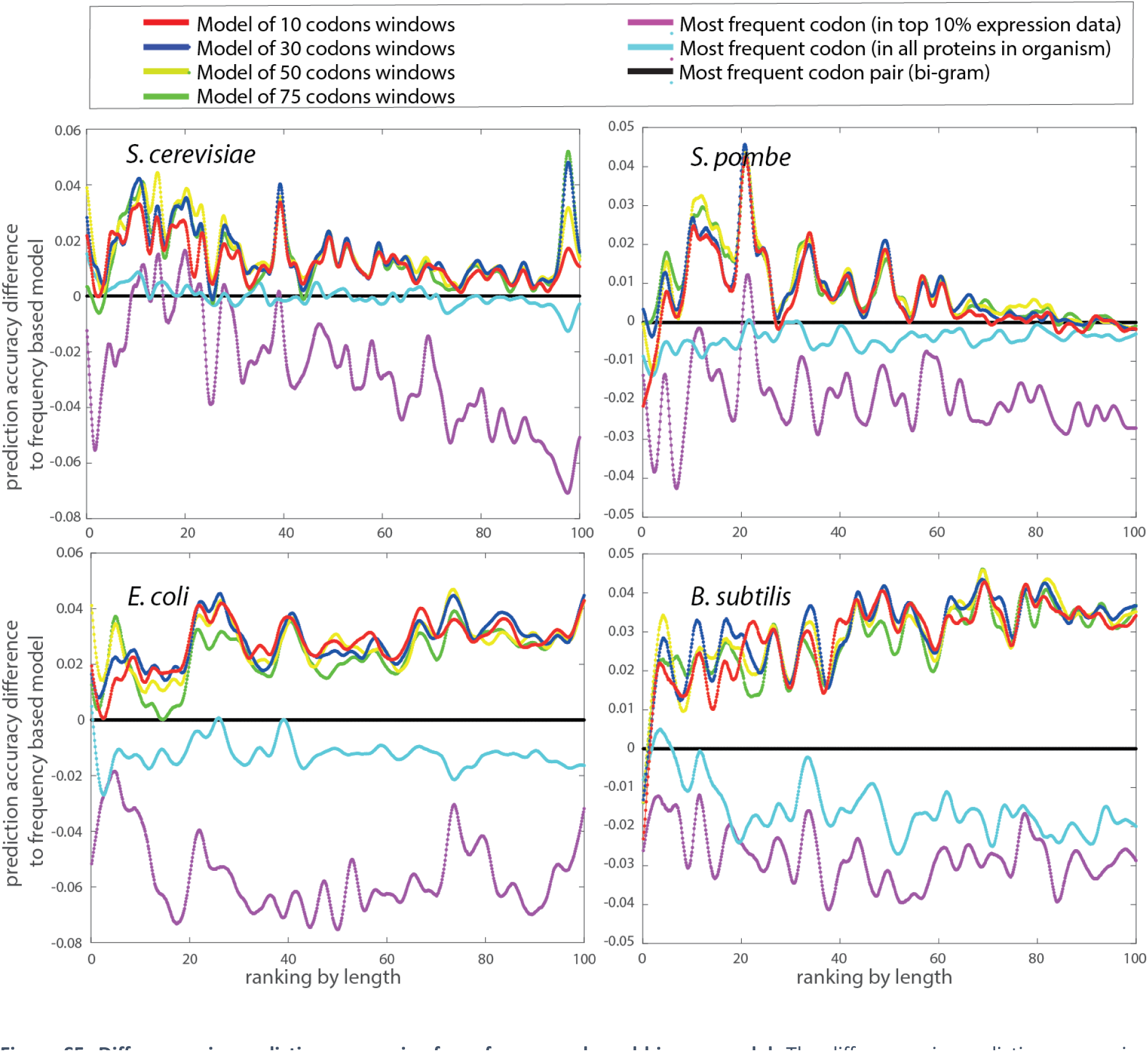
Differences in prediction accuracies from frequency-based bigram model. The differences in prediction accuracies between the mBART models and the frequency-based bigram model are plotted versus length rankings. These are the same data as in Figure 4. The frequency-based bigram model accuracies (black) are shown as a fixed value of 0. In this figure as well, we see that in all four organisms, the mBART models outperform the frequency-based baseline, and that the advantage is greater in the bacterial organisms.

**Figure S6:**
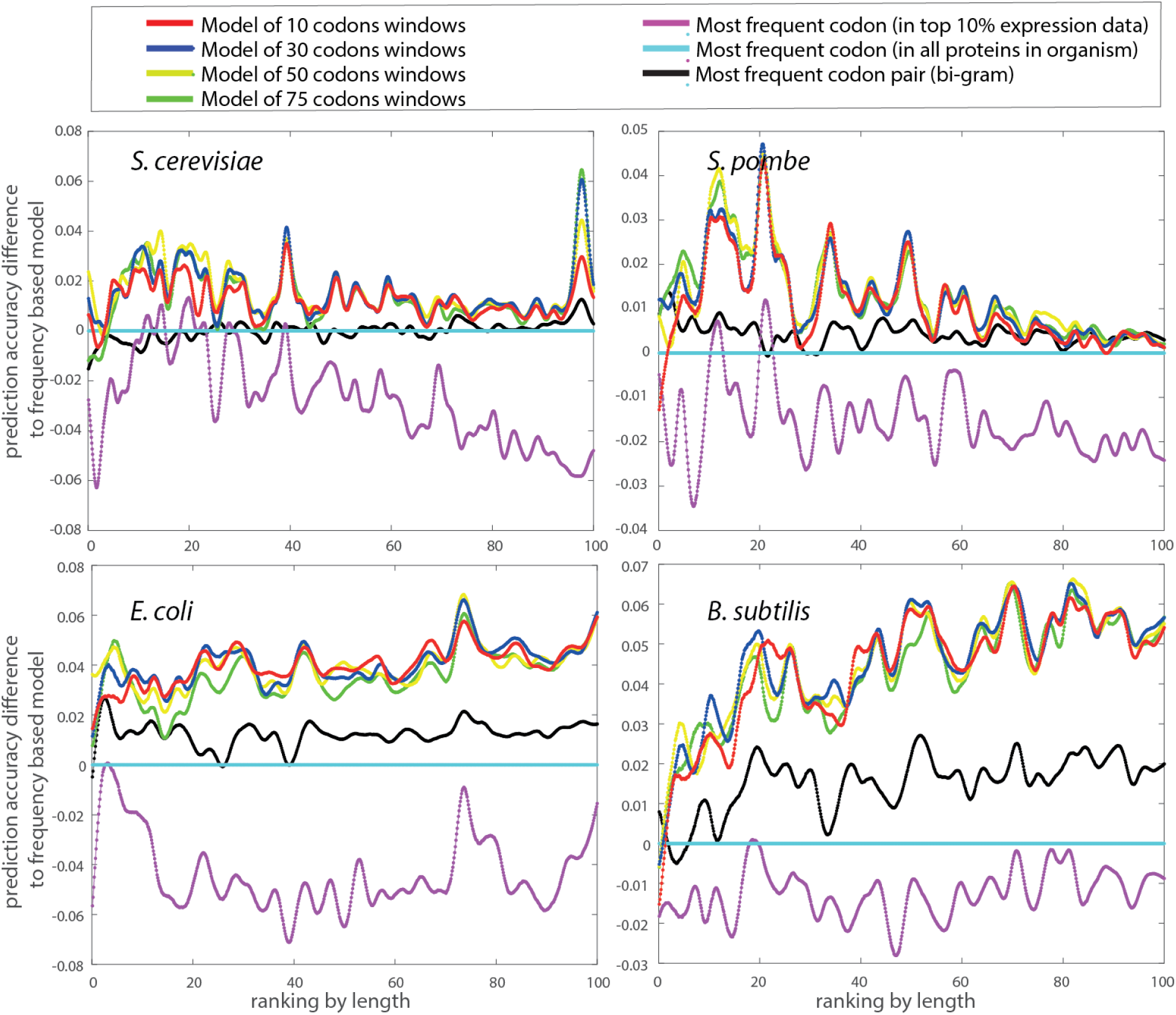
Codon prediction accuracies with inference in masking mode with respect to protein length. The differences in prediction accuracies between the mBART models and the frequency-based model plotted versus proteins ranked by length. These are the same data as in Figure 4, only the *y*-axis plots the differences in accuracies with respect to the frequency-based model. Thus, the cyan frequency-based baseline is shown as a fixed value of 0.

**Figure S7:**
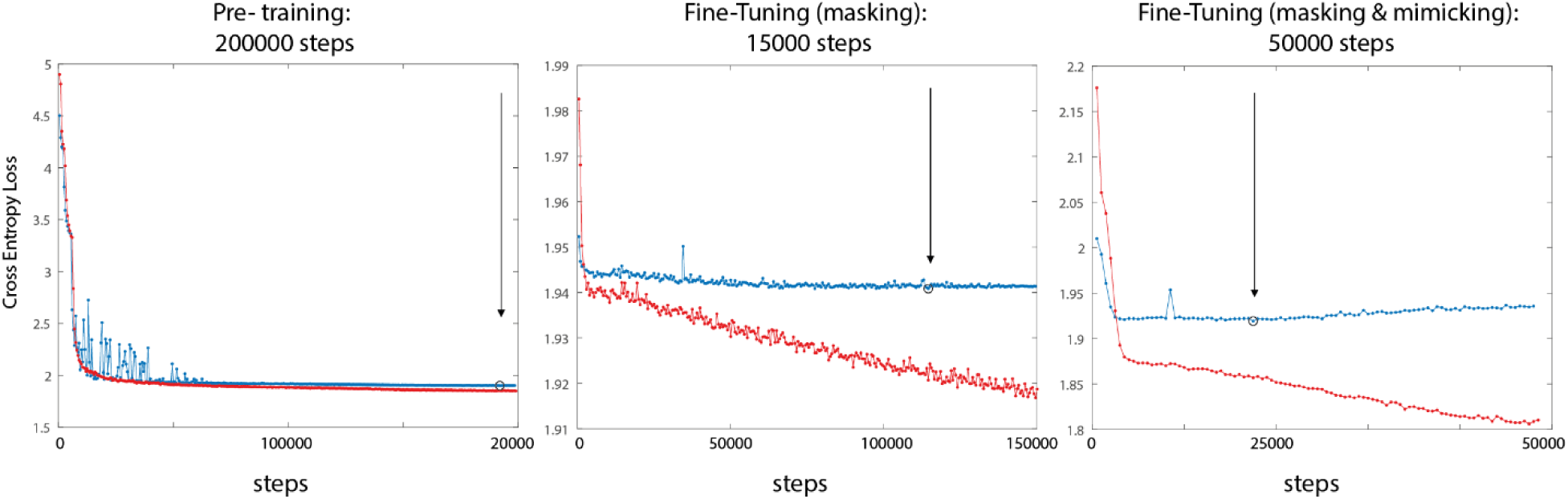
Cross entropy loss during training of the 30 codons window models. The training data was separated to train and validation sets, the loss on the training is shown in red, and the loss on the validation in blue. (Left) First, we pre-trained the model for 200,000 steps; the task is masking and mimicking, and the datasets support these tasks. Then, we select the model with the lowest loss on the validation set (marked by an arrow). (Middle) This model was then fine-tuned only on the masking task for 150,000 more steps (with the corresponding dataset). The model with the lowest loss on the validation set (marked by an arrow) was selected. (Right) In cases we wanted to use a model that was trained on mimicking, we further fine-tuned this model on the masking & mimicking tasks for 50,000 more steps. The model with the lowest loss on the validation set (marked by an arrow) was selected.

**Figure S8:**
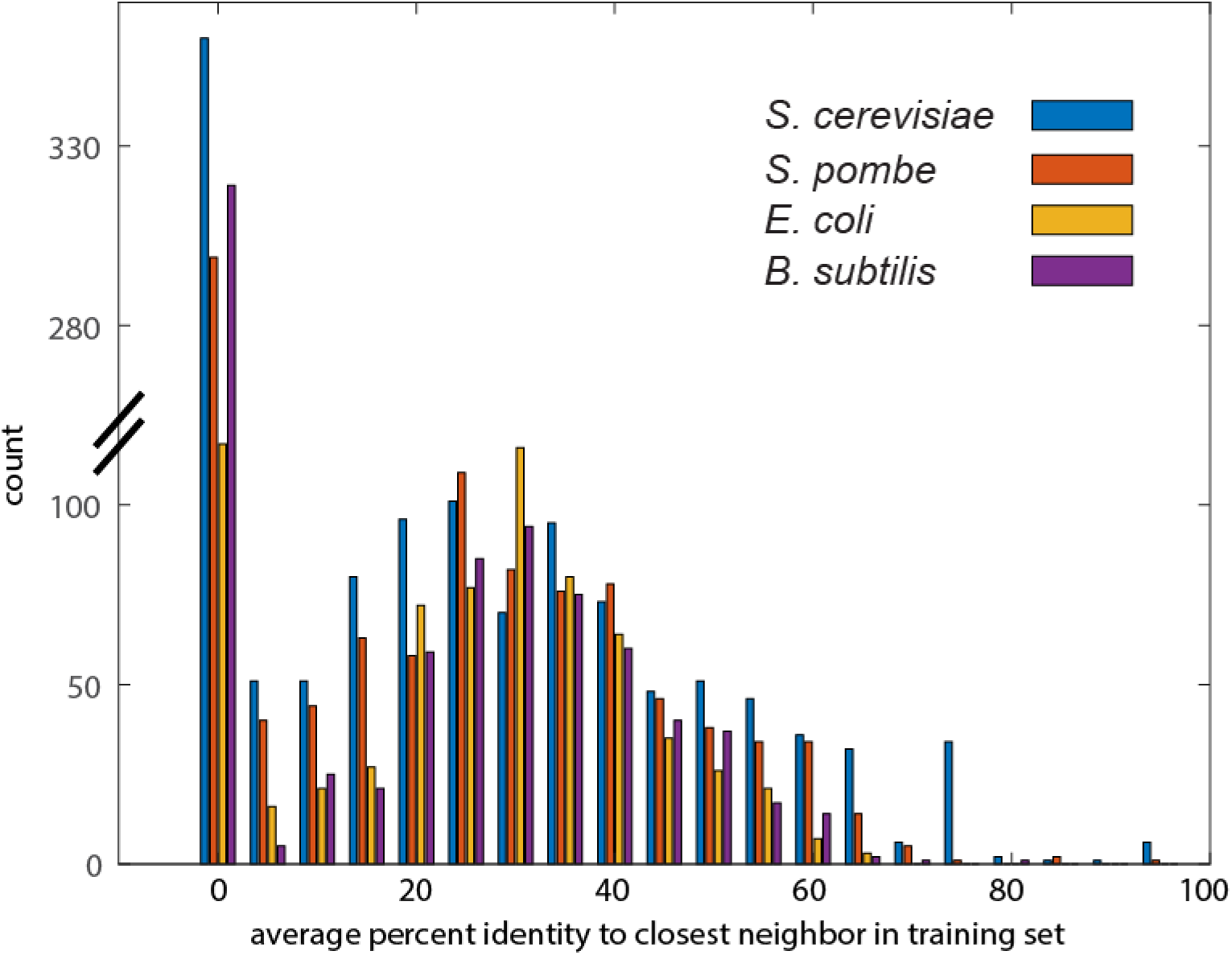
Cumulative prediction accuracies for the mimicking test set of aligned proteins with the proteins sorted by the accuracy of the prediction.

**Figure S9:**
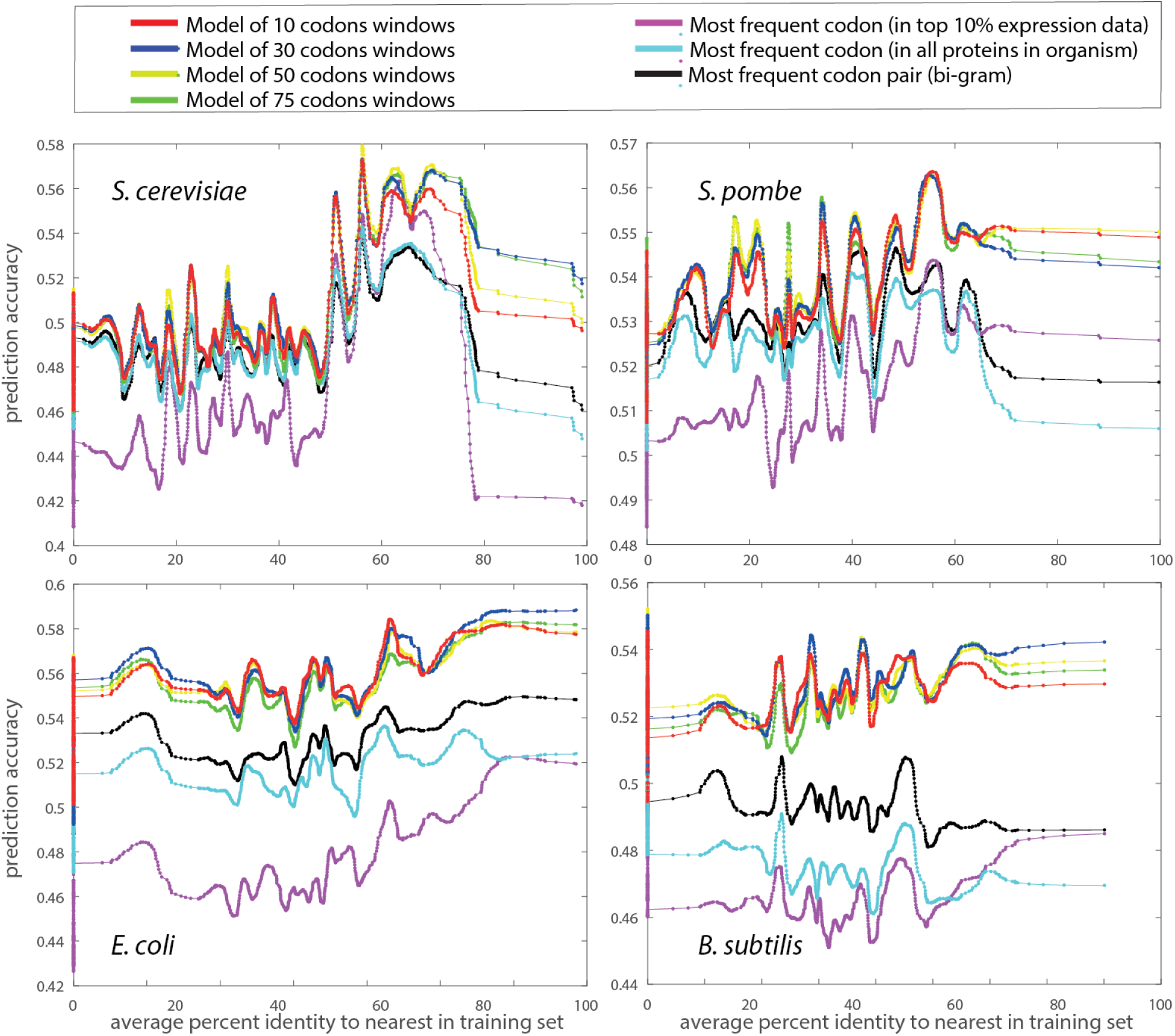
Codon prediction accuracies for the test set sorted by the average percent identity of the nearest protein in the training set. The values along the y-axis are the average accuracies of the set of proteins with the accuracy at that x-axis percent identity.

**Figure S10:**
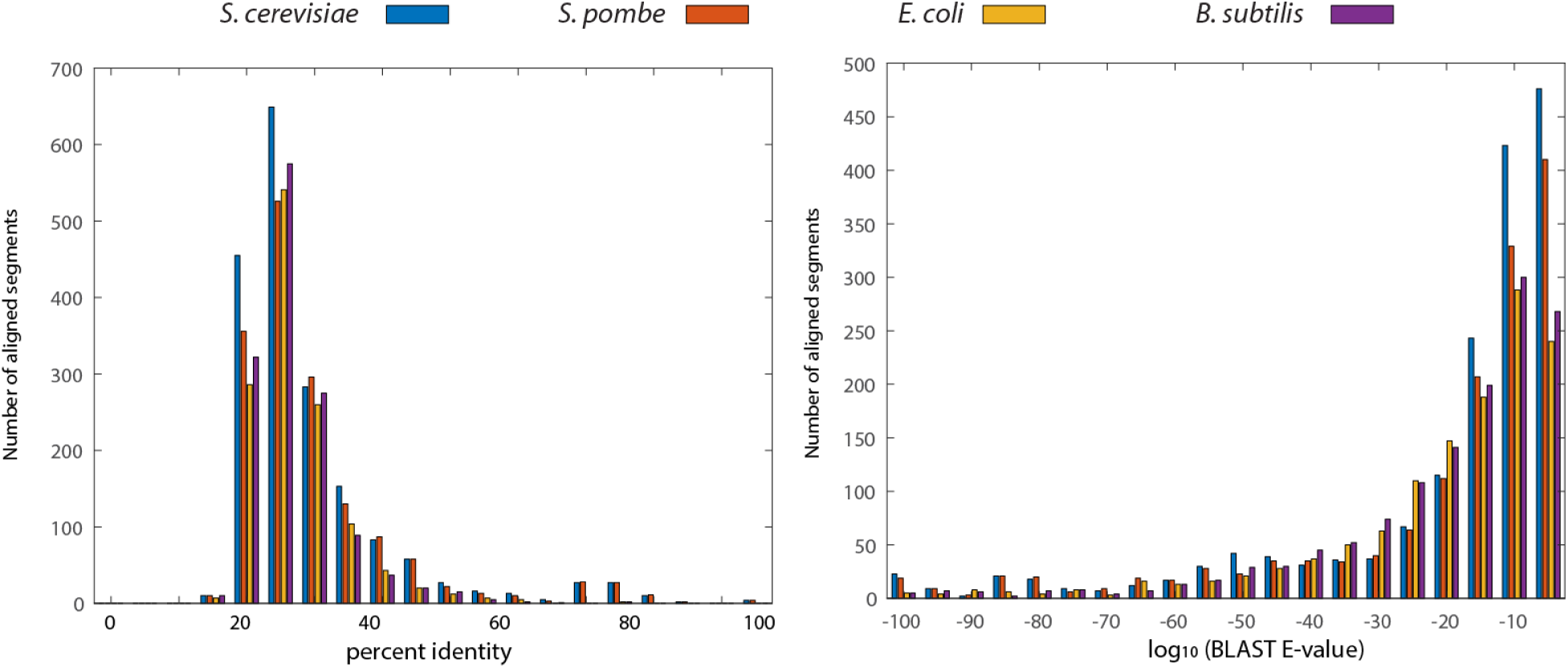
Histograms of the percent similarity and log(E-value) in the ortholog test set grouped by organism. The pairs of orthologs found by BLAST with an E-value less than 0.01 have low E-values, with many of them lower than 10^-10^, yet the percent identity is lower (the mean values are 32%, 33%, 30%, and 29% for *S. cerevisiae*, *S. pombe*, *E. coli*, and *B. subtilis*, respectively)

**Figure S11:**
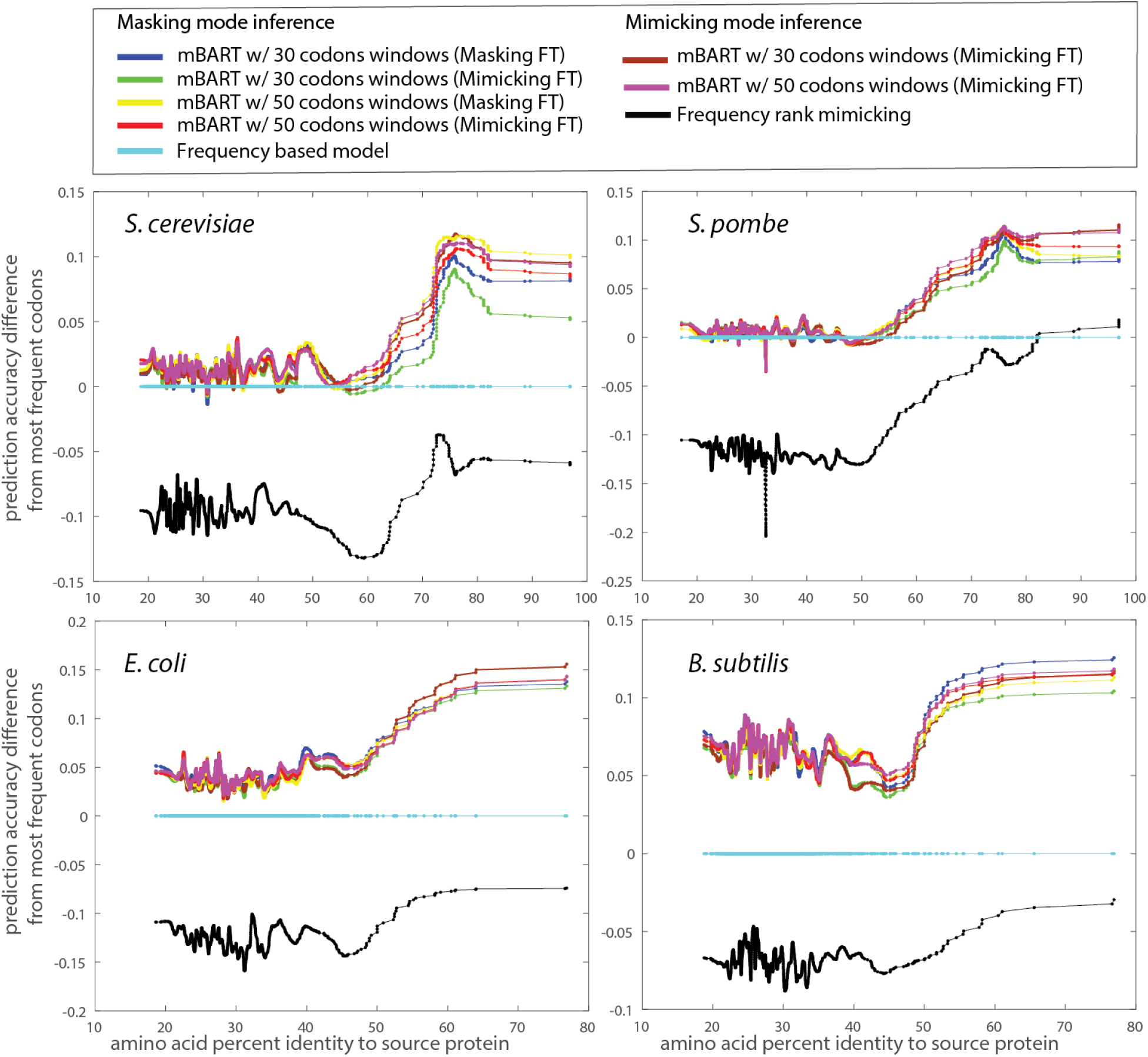
Codon prediction accuracies of mBART models in mimicking and masking modes compared to the frequency-based baseline with proteins sorted by the percent amino acid identity to their orthologף. This is the same data as in Figure 5, only the *y*-axis plots the differences in accuracies with respect to the frequency-based model. Thus, the cyan frequency-based baseline is shown as a fixed value of 0.

**Figure S13:**
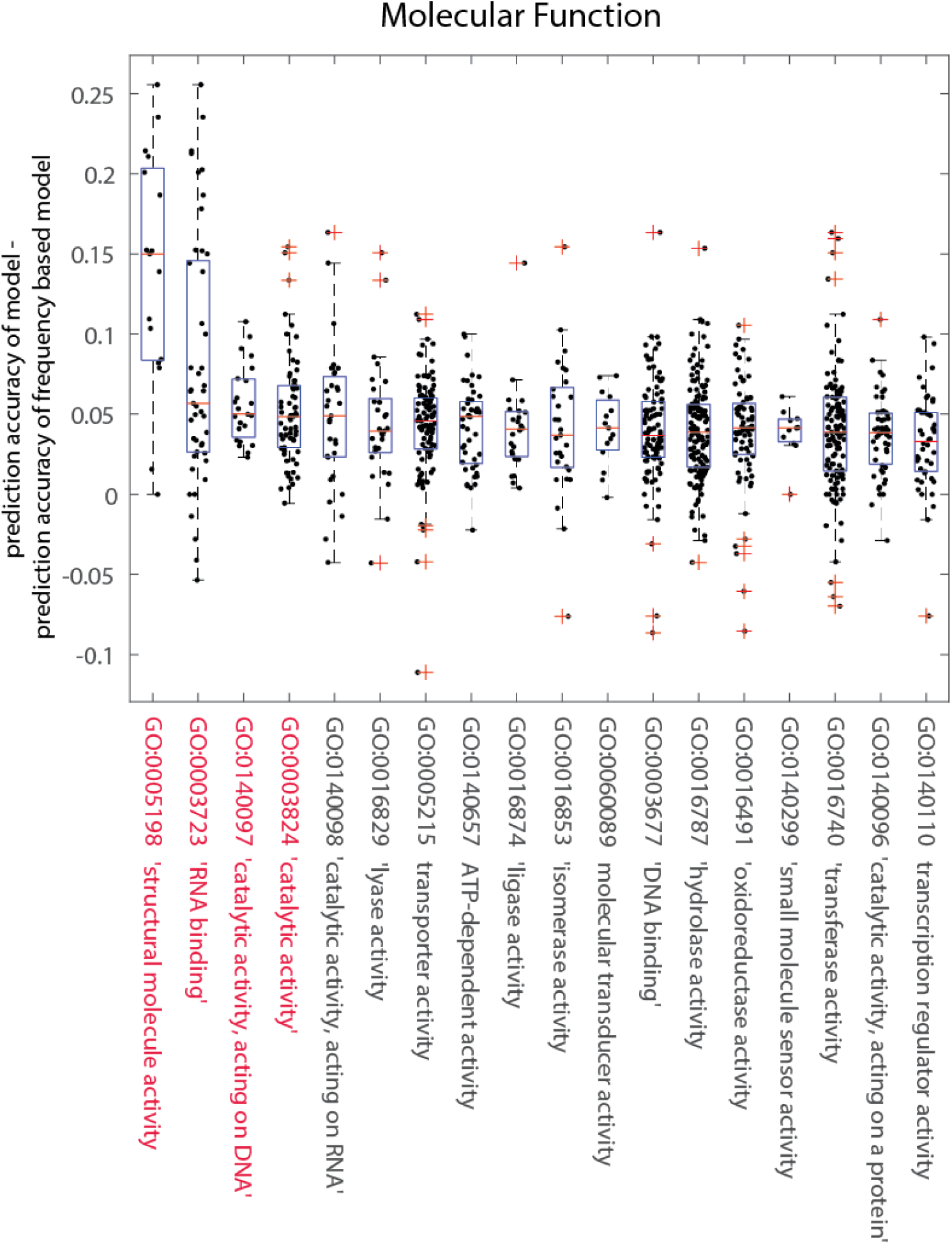
Prediction accuracy of *E. coli* test-set proteins grouped by GO molecular function. Differences between masking-task FT mBART model (30-codon window) predictions and the frequency-based baseline for test-set proteins grouped by GO terms. Terms were sorted in descending order by the mean average difference. The terms for which the p-value is <0.05 in a Mann-Whitney rank sum test for the functionally grouped set versus the rest of the test-set proteins are highlighted in red.

**Figure S14:**
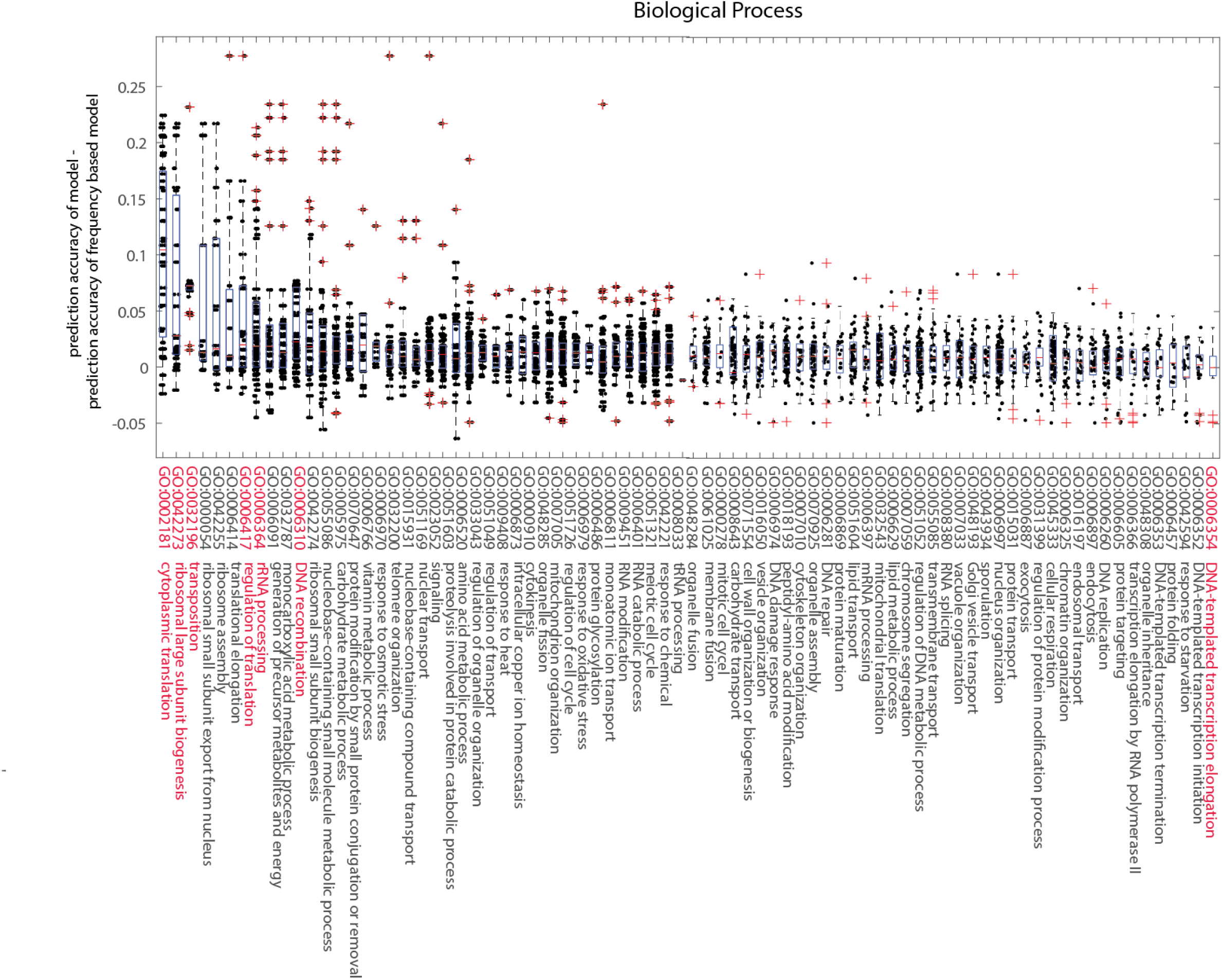
Prediction accuracy of *S. cerevisiae* test-set proteins grouped by GO biological process. Differences between masking-task FT mBART model (30-codon window) predictions and the frequency-based baseline for test-set proteins grouped by GO terms. Terms were sorted in descending order by the mean average difference. The terms for which the p-value is <0.05 in a Mann-Whitney rank sum test for the functionally grouped set versus the rest of the test-set proteins are highlighted in red.

**Figure S15:**
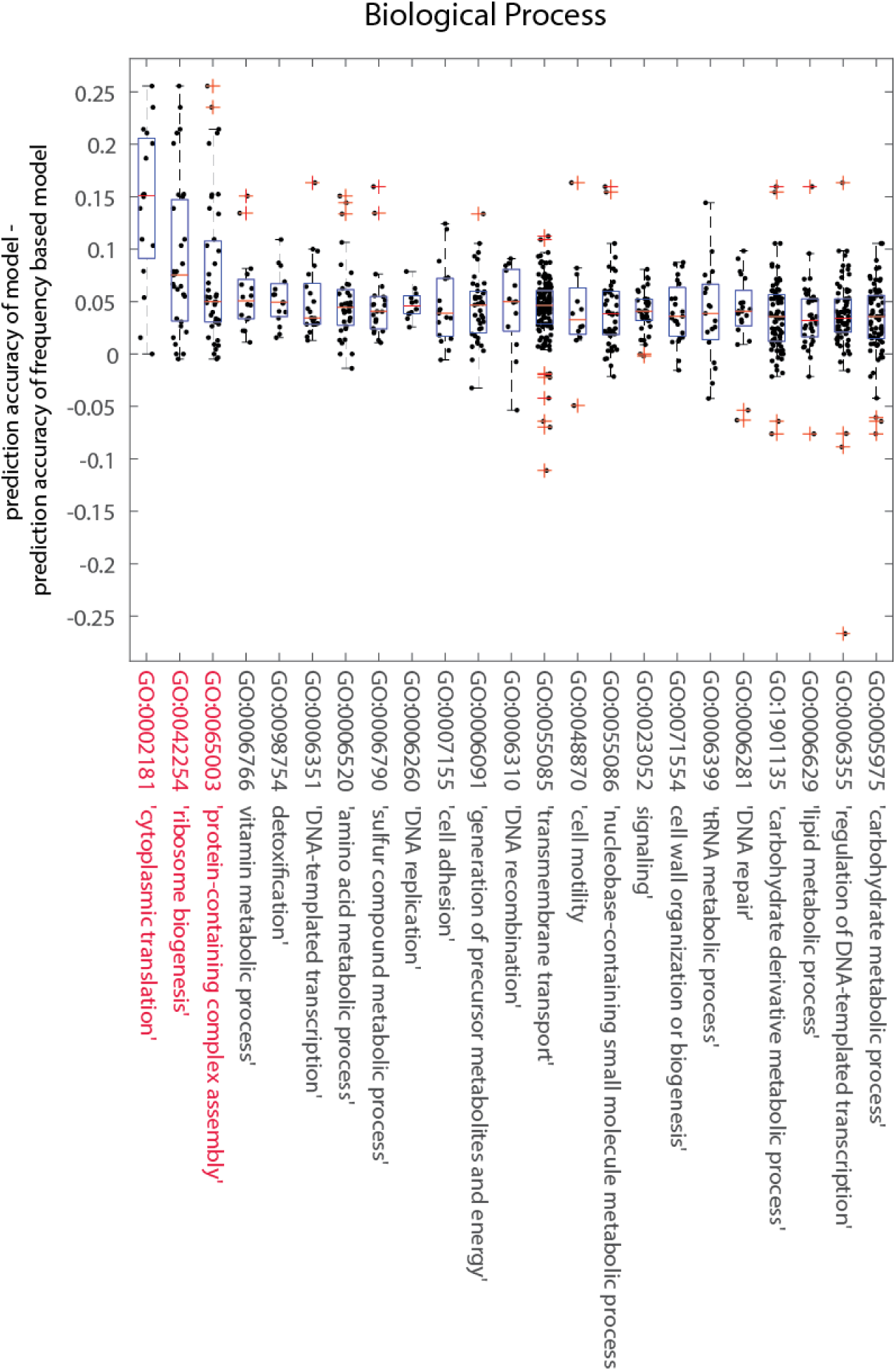
Prediction accuracy of *E. coli* test-set proteins grouped by GO biological process. Differences between masking-task FT mBART model (30-codon window) predictions and the frequency-based baseline for test-set proteins grouped by GO terms. Terms were sorted in descending order by the mean average difference. The terms for which the p-value is <0.05 in a Mann-Whitney rank sum test for the functionally grouped set versus the rest of the test-set proteins are highlighted in red.

**Figure S16:**
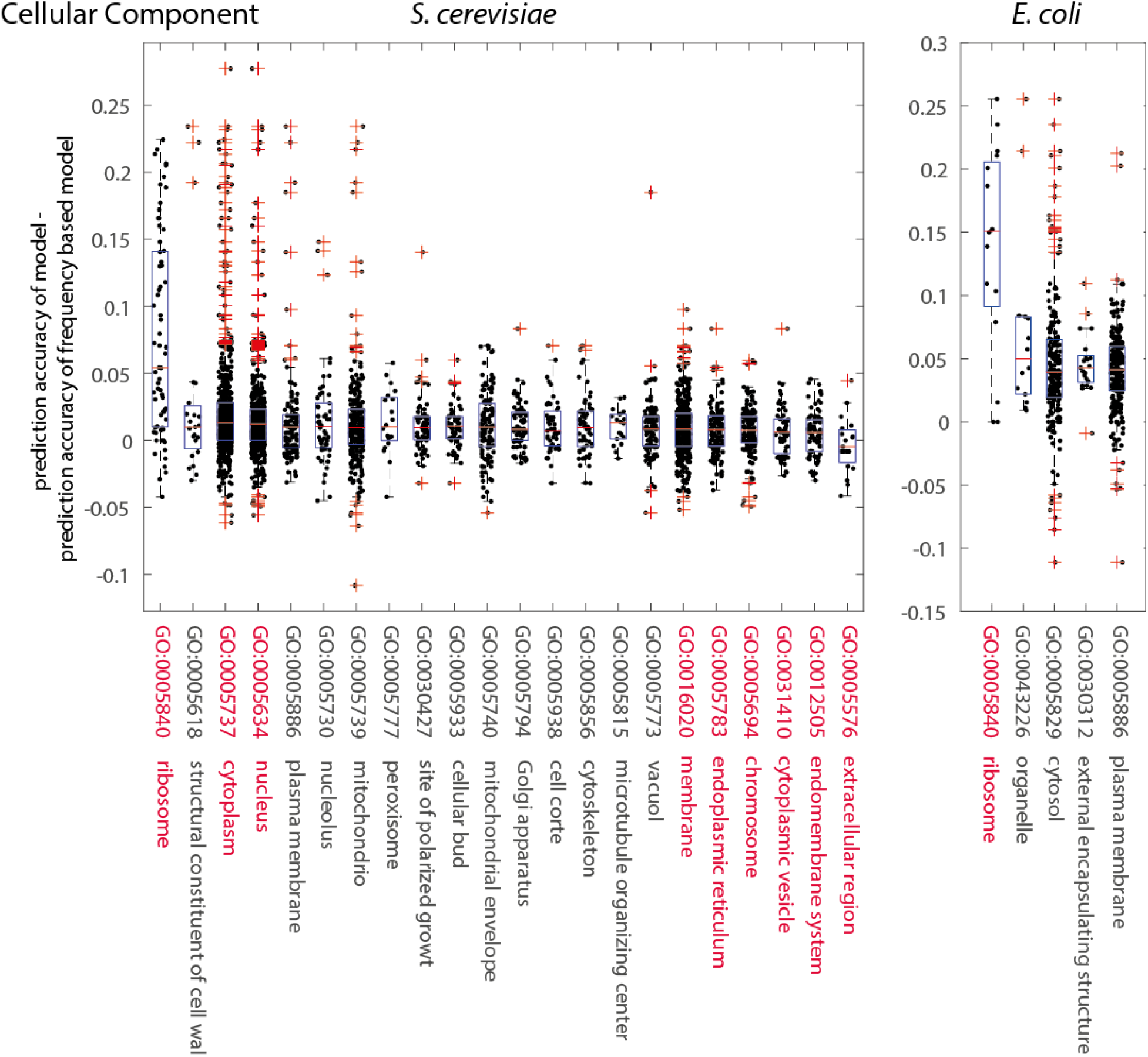
Prediction accuracy of *S. cerevisiae* and *E. coli* test-set proteins grouped by GO cellular component terms. Differences between masking-task FT mBART model (30-codon window) predictions and the frequency-based baseline for test-set proteins grouped by GO terms. Terms were sorted in descending order by the mean average difference. The terms for which the p-value is <0.05 in a Mann-Whitney rank sum test for the functionally grouped set versus the rest of the test-set proteins are highlighted in red.

**Table S1:** Significance of differences in accuracies between models. P-values and Normalize Cohen d values were determined using a paired-sample t-test on the test set proteins (grouped by organism) for the test decision for the null hypothesis that differences between models comes from a normal distribution with mean equal to zero and unknown variance. The 30 codons window is better than the other models, as evidenced by the absolute mean percentage difference, and p-values indicating that the difference is significant although the Cohen d value showing that the effect size is not big.

| Compared window sizes | *S. cerevisiae* |  |  | *S. pombe* |  |  |
| --- | --- | --- | --- | --- | --- | --- |
|  | p-value | effect size |  | p-value | effect size |  |
|  |  | Absolute (<% diff>) | Normalized (Cohen d) |  | Absolute (<% diff>) | Normalized (Cohen d) |
| 30 vs. 10 | **<10<sup>-5</sup>** | 0.2895 | 0.0448 | **0.0011** | 0.1531 | 0.0338 |
| 30 vs. 50 | 0.2514 | -0.0601 | -0.0031 | 0.0775 | -0.0750 | -0.0003 |
| 30 vs. 75 | **0.0078** | 0.1560 | 0.0278 | 0.1065 | -0.0752 | -0.0036 |
| 50 vs. 10 | **<10<sup>-5</sup>** | 0.3495 | 0.0489 | **<10<sup>-5</sup>** | 0.2280 | 0.0341 |
| 50 vs. 75 | **0.0002** | 0.2161 | 0.0315 | 0.9945 | -0.0003 | -0.0033 |
| 75 vs. 10 | 0.0567 | 0.1334 | 0.0159 | **0.0001** | 0.2283 | 0.0374 |
|  | *E. coli* |  |  | *B. subtilis* |  |  |
|  | p-value | effect size |  | p-value | effect size |  |
|  |  | Absolute (<% diff>) | Normalized (Cohen d) |  | Absolute (<% diff>) | Normalized (Cohen d) |
| 30 vs. 10 | 0.6065 | 0.0452 | -0.0185 | **0.0297** | 0.1766 | 0.0489 |
| 30 vs. 50 | 0.2584 | 0.0933 | 0.0149 | 0.4422 | 0.0608 | 0.0152 |
| 30 vs. 75 | **<10<sup>-5</sup>** | 0.4624 | 0.0731 | **0.0037** | 0.2576 | 0.0672 |
| 50 vs. 10 | 0.5722 | -0.0480 | -0.0338 | 0.2047 | 0.1158 | 0.0347 |
| 50 vs. 75 | **<10<sup>-5</sup>** | 0.3691 | 0.0600 | **0.0123** | 0.1968 | 0.0534 |
| 75 vs. 10 | **<10<sup>-5</sup>** | -0.4172 | -0.0914 | 0.4146 | -0.0810 | -0.0181 |

**Table S2:** Significance of differences in accuracies between models. The p-values were determined using a paired-sample *t*-test on the test set proteins (grouped by organism). The p-value is for the test decision for the null hypothesis that the differences between models comes from a normal distribution with mean equal to zero and unknown variance.

| Compared predictions | *S. cerevisiae* |  |  | *S. pombe* |  |  |
| --- | --- | --- | --- | --- | --- | --- |
|  | p-value | effect size |  | p-value | effect size |  |
|  |  | Absolute (<% diff>) | Normalized (Cohen d) |  | Absolute (<% diff>) | Normalized (Cohen d) |
| masking-FT-masking vs. mimicking-FT-masking | 1.41e-04 | 0.2119 | 0.2119 | 5.32e-01 | -0.0315 | -0.0315 |
| mimicking-FT-mimicking vs. mimicking-FT-masking | <10<sup>-5</sup> | 0.2125 | 0.2125 | 1.9394e-03 | 0.1255 | 0.1255 |
| mimicking-FT-mimicking vs. masking-FT-masking | 9.92e-01 | 0.0006 | 0.0006 | 5.83e-03 | 0.1570 | 0.1570 |
| mimicking-FT-mimicking vs. frequency-rank-mimicking | <10<sup>-5</sup> | 10.8005 | 10.8005 | <10<sup>-5</sup> | 12.3473 | 12.3473 |
| mimicking-FT-mimicking vs. frequency-bigram | <10<sup>-5</sup> | 1.4005 | 1.4005 | <10<sup>-5</sup> | 0.6559 | 0.6559 |
|  | *E. coli* |  |  | *B. subtilis* |  |  |
|  | p-value | effect size |  | p-value | effect size |  |
|  |  | Absolute (<% diff>) | Normalized (Cohen d) |  | Absolute (<% diff>) | Normalized (Cohen d) |
| masking-FT-masking vs. mimicking-FT-masking | <10<sup>-5</sup> | 0.3303 | 0.3303 | <10<sup>-5</sup> | 0.4589 | 0.4589 |
| mimicking-FT-mimicking vs. mimicking-FT-masking | 2.72e-01 | 0.0492 | 0.0492 | 2.28e-02 | 0.1068 | 0.1068 |
| mimicking-FT-mimicking vs. masking-FT-masking | 3.58e-05 | -0.2811 | -0.2811 | <10<sup>-5</sup> | -0.3521 | -0.3521 |
| mimicking-FT-mimicking vs. frequency-rank-mimicking | <10<sup>-5</sup> | 16.3095 | 16.3095 | <10<sup>-5</sup> | 13.2083 | 13.2083 |
| mimicking-FT-mimicking vs. frequency-bigram | <10<sup>-5</sup> | 2.2391 | 2.2391 | <10<sup>-5</sup> | 4.0171 | 4.0171 |

**Table S3A:** P-values in a Mann-Witney rank sum test comparing the accuracy difference values for S. cerevisiae proteins with a specific GO annotation versus the accuracy difference of all other proteins in the test-set. Those with p-values < 0.05 are marked in red.

| *S. cerevisiae* |  |  |  |  |  |  |  |
| --- | --- | --- | --- | --- | --- | --- | --- |
| Molecular Function |  | Biological Process |  |  |  | Cellular Component |  |
| GO | p-value<br>Mann<br>Witney<br>rank sum<br>test | GO | p-value<br>Mann<br>Witney<br>rank sum<br>test | GO | p-value<br>Mann Witney<br>rank sum test | GO | p-value<br>Mann Witney<br>rank sum test |
| GO:0003735 | <10<sup>-5</sup> | GO:0002181 | <10<sup>-5</sup> | GO:0061025 | 0.9506 | GO:0005840 | <10<sup>-5</sup> |
| GO:0019843 | 0.0090 | GO:0042273 | 0.0001 | GO:0000278 | 0.9723 | GO:0005618 | 0.9524 |
| GO:0003723 | <10<sup>-5</sup> | GO:0032196 | <10<sup>-5</sup> | GO:0008643 | 0.4526 | GO:0005737 | <10<sup>-5</sup> |
| GO:0016779 | <10<sup>-5</sup> | GO:0000054 | 0.1228 | GO:0071554 | 0.8885 | GO:0005634 | 0.0227 |
| GO:0004518 | <10<sup>-5</sup> | GO:0042255 | 0.0696 | GO:0016050 | 0.6158 | GO:0005886 | 0.3256 |
| GO:0008233 | <10<sup>-5</sup> | GO:0006414 | 0.2034 | GO:0006974 | 0.8160 | GO:0005730 | 0.8958 |
| GO:0005085 | 0.1016 | GO:0006417 | 0.0345 | GO:0018193 | 0.8294 | GO:0005739 | 0.6064 |
| GO:0008135 | 0.8778 | GO:0006364 | 0.0035 | GO:0007010 | 0.6400 | GO:0005777 | 0.6868 |
| GO:0003677 | <10<sup>-5</sup> | GO:0006091 | 0.1299 | GO:0070925 | 0.8545 | GO:0030427 | 0.8304 |
| GO:0003729 | 0.5039 | GO:0032787 | 0.0986 | GO:0006281 | 0.5732 | GO:0005933 | 0.9379 |
| GO:0016740 | <10<sup>-5</sup> | GO:0006310 | <10<sup>-5</sup> | GO:0006869 | 0.6647 | GO:0005740 | 0.8002 |
| GO:0019899 | 0.7018 | GO:0042274 | 0.2192 | GO:0051604 | 0.8311 | GO:0005794 | 0.8312 |
| GO:0016491 | 0.8055 | GO:0055086 | 0.2457 | GO:0006397 | 0.3359 | GO:0005938 | 0.5762 |
| GO:0043167 | 0.0857 | GO:0005975 | 0.0835 | GO:0032543 | 0.7184 | GO:0005856 | 0.4681 |
| GO:0016301 | 0.0680 | GO:0070647 | 0.0893 | GO:0006629 | 0.4877 | GO:0005815 | 0.9855 |
| GO:0003682 | 0.8091 | GO:0006766 | 0.3234 | GO:0007059 | 0.3712 | GO:0005773 | 0.1235 |
| GO:0016787 | 0.5941 | GO:0006970 | 0.2120 | GO:0051052 | 0.4888 | GO:0016020 | 0.0034 |
| GO:0016829 | 0.8789 | GO:0032200 | 0.6492 | GO:0055085 | 0.2004 | GO:0005783 | 0.0196 |
| GO:0030674 | 0.6009 | GO:0015931 | 0.6253 | GO:0008380 | 0.5192 | GO:0005694 | 0.0387 |
| GO:0030234 | 0.3608 | GO:0051169 | 0.8188 | GO:0007033 | 0.6177 | GO:0031410 | 0.0385 |
| GO:0016853 | 0.9891 | GO:0023052 | 0.2371 | GO:0048193 | 0.1997 | GO:0012505 | 0.0095 |
| GO:0016874 | 0.8552 | GO:0051603 | 0.8518 | GO:0043934 | 0.3572 | GO:0005576 | 0.0037 |
| GO:0140657 | 0.3864 | GO:0006520 | 0.9124 | GO:0006997 | 0.4709 |  |  |
| GO:0003700 | 0.4078 | GO:0033043 | 0.9074 | GO:0015031 | 0.0528 |  |  |
| GO:0016757 | 0.9401 | GO:0051049 | 0.4584 | GO:0006887 | 0.3451 |  |  |
| GO:0008289 | 0.0596 | GO:0009408 | 0.9932 | GO:0031399 | 0.6340 |  |  |
| GO:0005198 | 0.6686 | GO:0006873 | 0.7386 | GO:0045333 | 0.3696 |  |  |
| GO:0022857 | 0.1204 | GO:0000910 | 0.8651 | GO:0006325 | 0.1272 |  |  |
| GO:0008168 | 0.8210 | GO:0048285 | 0.8565 | GO:0016197 | 0.1116 |  |  |
| GO:0016798 | 0.7158 | GO:0007005 | 0.4369 | GO:0006897 | 0.0512 |  |  |
| GO:0042393 | 0.7465 | GO:0051726 | 0.2588 | GO:0006260 | 0.1323 |  |  |
| GO:0016791 | 0.3677 | GO:0006979 | 0.5652 | GO:0006605 | 0.1023 |  |  |
| GO:0008092 | 0.0799 | GO:0006486 | 0.7698 | GO:0006366 | 0.1205 |  |  |
| GO:0032182 | 0.0751 | GO:0006811 | 0.1357 | GO:0048308 | 0.1877 |  |  |
| GO:0003924 | 0.0762 | GO:0009451 | 0.9211 | GO:0006353 | 0.0540 |  |  |
| GO:0051082 | 0.2990 | GO:0006401 | 0.9705 | GO:0006457 | 0.1107 |  |  |
| GO:0008134 | 0.1088 | GO:0051321 | 0.8960 | GO:0042594 | 0.0520 |  |  |
|  |  | GO:0042221 | 0.6498 | GO:0006352 | 0.0698 |  |  |
|  |  | GO:0008033 | 0.8562 | GO:0006354 | 0.0263 |  |  |
|  |  | GO:0048284 | 0.9129 |  |  |  |  |

**Table S3B:** P-values in a Mann-Witney rank sum test comparing the accuracy difference values for E. coli proteins with a specific GO annotation versus the accuracy difference of all other proteins in the test-set. Those with p-values < 0.05 are marked in red.

| E. coli |  |  |  |  |  |
| --- | --- | --- | --- | --- | --- |
| Molecular Function |  | Biological Process |  | Cellular Component |  |
| GO | p-value<br>Mann Witney rank<br>sum test | GO | p-value<br>Mann Witney rank<br>sum test | GO | p-value<br>Mann Witney rank<br>sum test |
| GO:0005198 | <10<sup>-5</sup> | GO:0002181 | <10<sup>-5</sup> | GO:0005840 | <10<sup>-5</sup> |
| GO:0003723 | 0.0015 | GO:0042254 | 0.0001 | GO:0043226 | 0.1881 |
| GO:0140097 | 0.0160 | GO:0065003 | 0.0048 | GO:0005694 | 0.5024 |
| GO:0003824 | 0.0210 | GO:0071941 | 0.1022 | GO:0005829 | 0.4081 |
| GO:0140098 | 0.2749 | GO:0006766 | 0.1159 | GO:0030312 | 0.6730 |
| GO:0016829 | 0.7391 | GO:0006575 | 0.4122 |  |  |
| GO:0005215 | 0.0742 | GO:0016071 | 0.3001 |  |  |
| GO:0140657 | 0.6338 | GO:0098754 | 0.6134 |  |  |
| GO:0016874 | 0.7652 | GO:0030163 | 0.3152 |  |  |
| GO:0016853 | 0.9730 | GO:0006351 | 0.7923 |  |  |
| GO:0060089 | 0.8957 | GO:0098542 | 0.5450 |  |  |
| GO:0003677 | 0.8384 | GO:0006520 | 0.3783 |  |  |
| GO:0016787 | 0.6177 | GO:0006790 | 0.0388 |  |  |
| GO:0016491 | 0.6776 | GO:0006260 | 0.8224 |  |  |
| GO:0140299 | 0.9222 | GO:0007155 | 0.7543 |  |  |
| GO:0016740 | 0.5505 | GO:0061024 | 0.8530 |  |  |
| GO:0140096 | 0.3792 | GO:0006091 | 0.9267 |  |  |
| GO:0140110 | 0.1112 | GO:0006310 | 0.8524 |  |  |
|  |  | GO:0055085 | 0.8830 |  |  |
|  |  | GO:0048870 | 0.1467 |  |  |
|  |  | GO:0055086 | 0.2714 |  |  |
|  |  | GO:0023052 | 0.2496 |  |  |
|  |  | GO:0071554 | 0.1422 |  |  |

